# Individual small in-stream barriers contribute little to strong local population genetic structure five strictly aquatic macroinvertebrate taxa

**DOI:** 10.1101/2020.10.01.322222

**Authors:** Martina Weiss, Hannah Weigand, Florian Leese

## Abstract

Water flow in river networks is frequently regulated by man-made in-stream barriers. These obstacles can hinder dispersal of aquatic organisms and isolate populations leading to the loss of genetic diversity. Although millions of small in-stream barriers exist worldwide, such as weirs below 2 m or tunnels and pipes, their impact on the dispersal of macroinvertebrates with strictly aquatic life cycles is unclear. We therefore assessed the effects of such barriers on the population structure and effective dispersal of five macroinvertebrate species with strictly aquatic life cycles: the amphipod crustacean *Gammarus fossarum* (clade 11), three snail species of the *Ancylus fluviatilis* species complex and the flatworm *Dugesia gonocephala*. We studied populations at nine weirs and eight culverts (3 pipes, 5 tunnels), built 33-109 years ago, in the heavily fragmented catchment of the river Ruhr (Sauerland, Germany). To assess fragmentation and barrier effects, we generated genome-wide SNP data using ddRAD sequencing and evaluated clustering, differentiation between populations up- and downstream of each barrier and effective migration rates among sites and across barriers. Additionally, we applied population genomic simulations to assess expected differentiation patterns under different gene flow scenarios. Our data show that populations of all species are highly isolated at regional and local scales within few kilometres. While the regional population structure likely results from historical processes, the strong local differentiation suggests that contemporary dispersal barriers exist. However, we identified significant barrier effects only for pipes (for *A. fluviatilis* II and III) and few larger weirs (>1.3 m; for *D. gonocephala*). Therefore our data suggest that most small in-stream barriers can probably be overcome by all studied taxa frequently enough to prevent fragmentation. Thus, the barriers leading to population isolation after few kilometres still need to be identified. Specifically, it remains to be tested if the strong local differentiation is a result of a cumulative effect of small barriers, or if e.g. larger in-stream barriers, land use, chemical pollution, urbanisation, or a combination of these factors impede gene flow.

## Introduction

Rivers in the Anthropocene are in crisis due to various human activities (Jackson et al., 2001; Vörösmarty et al., 2010). In particular, natural water flow has been altered substantially over the past few centuries by the construction of in-stream barriers, such as dams, weirs (Grill et al., 2015; Nilsson et al., 2005; Zarfl et al., 2015) and culverts (David et al., 2014; Torterotot et al., 2014; Wheeler et al., 2005). Such human-induced habitat fragmentation is considered a major threat to biodiversity (Hudman & Gido, 2013). In isolated populations, the effects of genetic drift and inbreeding are higher, thereby increasing the risk of genetic diversity loss (e.g., Vrijenhoek, 1998) and extinction (Bijlsma & Loeschcke, 2012). To predict population resilience and long-term adaptability, it is therefore crucial to understand species-specific dispersal patterns (Hughes et al., 2008) and to identify actual barriers to dispersal.

Even within species, dispersal rates can differ between areas and sites as site-specific environmental factors such as water chemistry, land use and urbanisation can influence effective dispersal rates making predictions difficult. Another important factor that influences population genetic structure are in-stream barriers. Impacts of such barriers have been shown mainly for populations of different fish species (e.g., Hansen et al., 2014; Horreo et al., 2011; Junker et al., 2012; Torterotot et al., 2014; Van Leeuwen et al., 2018). In contrast, for freshwater invertebrate taxa, the effect of in-stream barriers is often unknown. Although barrier effects have been reported (David et al., 2014; Dillon, 1988; Resh, 2005), they are highly species-specific (Tonkin et al., 2014) owing to divergence in life history traits. While more studies have focused on large barriers such as dams or reservoirs, little is known about the effects of smaller obstacles such as weirs and culverts that can occur at high densities, with less than 1 km between barriers (Belletti et al., 2020; Dumont et al., 2005). Only limited information about the properties of these small barriers exists, especially with respect to barrier age, shape and impoundment areas, making it difficult to predict their impact on genetic structure. Further, in-stream barriers might not act as complete barriers, i.e. they may not prevent gene flow equally in both directions, but only, or stronger, in upstream direction. This asymmetry makes it more difficult to detect effects. When focusing on macroinvertebrates, other confounding factors that limit the detectability of barrier effects are the often large populations sizes (limiting genetic drift effects) and low dispersal abilities (e.g., in strictly aquatic taxa). Low effective dispersal alone can generate a strong genetic background structure, which needs to be accounted for when studying impacts of in-stream barriers on genetic structure (Coleman et al., 2018). Therefore, beside studying the direct effect of barriers, it is also important to know the general population structure of the species in the area to be able to interpret the results. In addition to temporal sampling or space-for-time study designs to distinguish between background genetic structure and barrier effects (Coleman et al., 2018), population genetic simulations can improve the interpretation of empirical data by generating null models under specific assumptions, for example life history traits (Hoban et al., 2012). Considering the relatively young age of many man-made barriers (typically tens of years to a few hundred years) and the often large population sizes of macroinvertebrate species, high density genomic markers, like single nucleotide polymorphisms (SNPs), are needed to resolve fine-scale patterns of genetic structure (Whiterod et al., 2017).

In this study, we first performed comparative population genomic analyses to assess effective dispersal of different and locally abundant stream macroinvertebrate taxa in a heavily fragmented German low mountain range stream network. The second aim was then to evaluate the impact of small in-stream barriers on population genetic structure in these species. To achieve these aims, we generated genome-wide SNP data by double digest restriction site-associated sequencing (ddRAD-seq, Peterson et al., 2012). To account for species- and site-specific genetic background structure, reference sites were sampled in each stream. Further, the distances between sites were chosen within the dispersal ability of all species (~200 m), so that without a barrier, gene flow should be high. All selected species, i.e. the amphipod crustacean *Gammarus fossarum* clade 11 (after Weiss et al., 2014), the flatworm *Dugesia gonocephala* and three snail species of the *Ancylus fluviatilis* species complex (*A. fluviatilis* I, II, III; Weiss et al., 2018), are strictly confined to the aquatic environment (i.e., are hololimnic) and commonly occur in small European streams. They differ in feeding type, reproductive strategy and dispersal mode. Based on specific information on ecology, life history traits (obtained from Baršiene et al., 1996; Brittain & Eikeland, 1988; Burch, 1962; de Vries, 1986; Eder et al., 1995; Elliott, 2003; MacNeil et al., 1997; Minshall & Winger, 1968; Moog, 1995; Nesemann & Reischütz, 1995; Patrick et al., 2014; Schmidt-Kloiber & Hering, 2015) and initial genetic data for *G. fossarum* (Alp et al., 2012; Weiss & Leese, 2016), we derived predictions concerning the strength of in-stream barrier effects and population structure (Figure 1). In order to put results into context, we simulated population genomic data as null models to test how species-specific life history traits influence the detectability of barrier effects. Based on mobility and population size (Figure 1), we hypothesised that *G. fossarum* shows the lowest and *A. fluviatilis* I, II and III the strongest population structure, while it is intermediate for *D. gonocephala*. Further, concerning barrier effects, we hypothesised that barriers increase genetic differentiation compared to control sites and that weirs have stronger effects than culverts. Additionally, we hypothesised that the effect of weirs depends on barrier properties, with the highest, steepest and smoothest barriers having the strongest effect. For culverts, we hypothesised that pipes have stronger effects than tunnels and that the effect depends on the length, diameter and bottom substrate of the culvert, with longer culverts with a smaller diameter and concrete bottom (instead of natural sediment) having the strongest effect. Furthermore, we hypothesise that effects of barriers on differentiation is species-specific, with strongest effects expected for *A. fluviatilis* I, II and III and weakest for *G. fossarum.*

**Figure 1.**
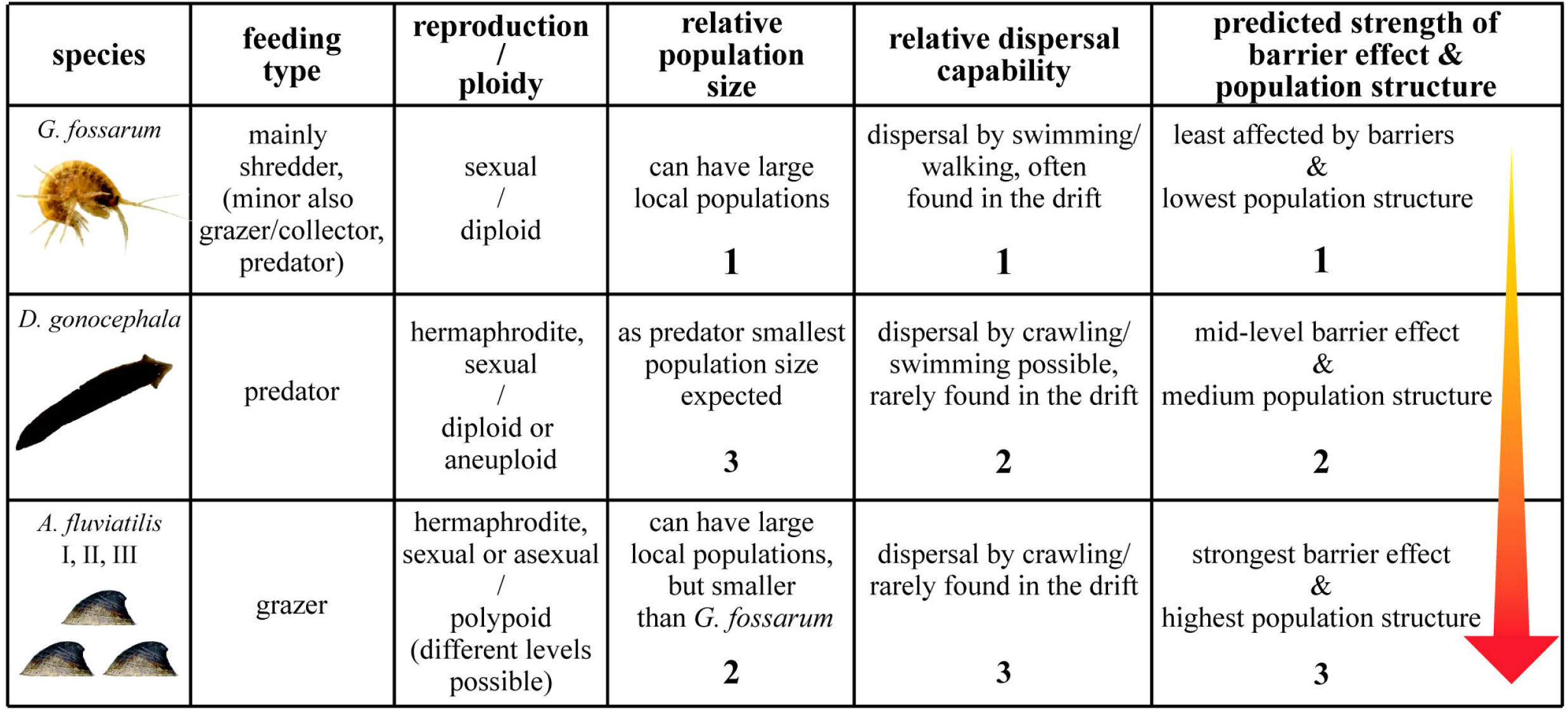
Summary of traits differing among studied species. Numbers from 1 to 3 indicate the expected relative gradient. The arrow indicates the expected strength of the barrier effects and population structure.

## Materials and Methods

### Study design and study sites

The study region was the river Ruhr catchment (North Rhine-Westphalia, Sauerland, Germany), with many weirs and culverts, often separated by less than 1 km (Dumont et al., 2005). Taxa sampled were *G. fossarum* clade 11, *D. gonocephala* and *A. fluviatilis.* The three species *A. fluviatilis* I, II and III can only be identified by ddRAD analyses (Weiss et al., 2018); accordingly, specimens were sampled and processed in the lab as a single species. The sampling scheme was designed to systematically test the impact of two types of in-stream barriers, weirs (abbreviated QB [for German ‘Querbauwerk’]) and culverts (VR [for German ‘Verrohrung’]), on gene flow. To compare gene flow among populations with and without a barrier in the same stream, specimens were sampled from at least three sampling sites (S), so that two sites were sampled upstream (S1 and S2) or downstream (S3 and S4) of the barrier with no barrier in-between. Where possible, reference sites were sampled up- and downstream of the barrier. In one case (VR17), three reference sites were sampled upstream of the barrier. If tributaries joined the stream within the sampling range, specimens were collected at these locations (abbreviated N). In two cases, the scheme was extended to contain five sites (S0–S4) with a barrier downstream of S1 and S2 (VR6 and QB27). Based on the dispersal range of all target taxa, 200 m was chosen as the distance between sampling sites, including sites separated by a barrier. The sampling stretch containing all individual sampling sites associated with one barrier is in the following called ‘barrier site’ (see Figure 2 and 3). Specimens were collected at 15 barrier sites consisting of 17 barriers (Figure 2), including eight culverts (8–120 m in length) and nine weirs (0.6–1.65 m in height) (Table 1, Figure S1 for exemplary pictures). At individual sampling sites, 5–12 specimens per species were sampled, but not all target species occurred at all sites (see Table S1 for details).

**Table 1.** Barrier characteristics of culverts (VR) and weirs (QB), sorted by expected severity from high to low. First three VRs were categorized as pipes, all other were tunnels.

| Name<br>VR/QB | Approx.<br>construction<br>year | Approx.<br>length VR/<br>height QB [m] | Approx.<br>width ×<br>height VR<br>[m] | VR bottom<br>substrate/ QB<br>quality<br>structure | Description of VR/<br>steepness of QB ramps<br>(low/med/high) |
| --- | --- | --- | --- | --- | --- |
| **VR20** | 1906<br>(enlarged 1980) | 94/114 | tunnel: 2.5 ×<br>1.2 pipe: ø<br>0.6 | concrete | tunnel (94 m) between N2<br>and S3 with a steep drop; S2<br>joins tunnel via ~30 m-long<br>pipe over small ramp<br>upstream of drop; total length<br>S1–S3: 114 m |
| **VR6b** | 1960 | 22 | ø 0.5 | concrete | 3 parallel concrete pipes,<br>small drop at end |
| **VR6a** | 1960 | 8 | ø 0.5 | concrete | 2 parallel concrete pipes,<br>small drop at end |
| **VR17** | 1980 | 120 | 3 × 1 | artificial<br>sediment | broad tunnel, small drops<br>between S1 & S2 |
| **VR23** | 1910<br>(enlarged 1970) | 70 | 0.6 × 1 | natural<br>sediment | tunnel |
| **VR12** | – | 38 | 1 × 1.2 | concrete | tunnel |
| **VR11** | 1970<br>(enlarged 1980) | 34 (72) | 3 × 2 | concrete | big tunnel, up- & downstream<br>7 small drops |
| **VR9** | 1980 | 56 | 3 × 4 | natural<br>sediment | big tunnel |
| **QB27a** | 1930 | 1.65 |  | smooth | high |
| **QB24** | 1930 | 1.5 |  | rough | high |
| **QB11** | 1930 | 1.35 |  | rough | medium, drops at start and<br>end |
| **QB12** | – | 1.1 |  | rough | 1st half medium, 2nd high |
| **QB27b** | 1930 | 1.05 |  | smooth | low, drops at start and end |
| **QB22** | 1905 | 1.3 |  | rough with<br>bigger stones | medium |
| **QB23** | 1930 | 1.2 |  | rough with<br>bigger stones | medium |
| **QB17** | 1960 | 0.6 |  | rough | medium |
| **QB20** | – | 1 |  | rough | low, small drop at end |

**Figure 2.**
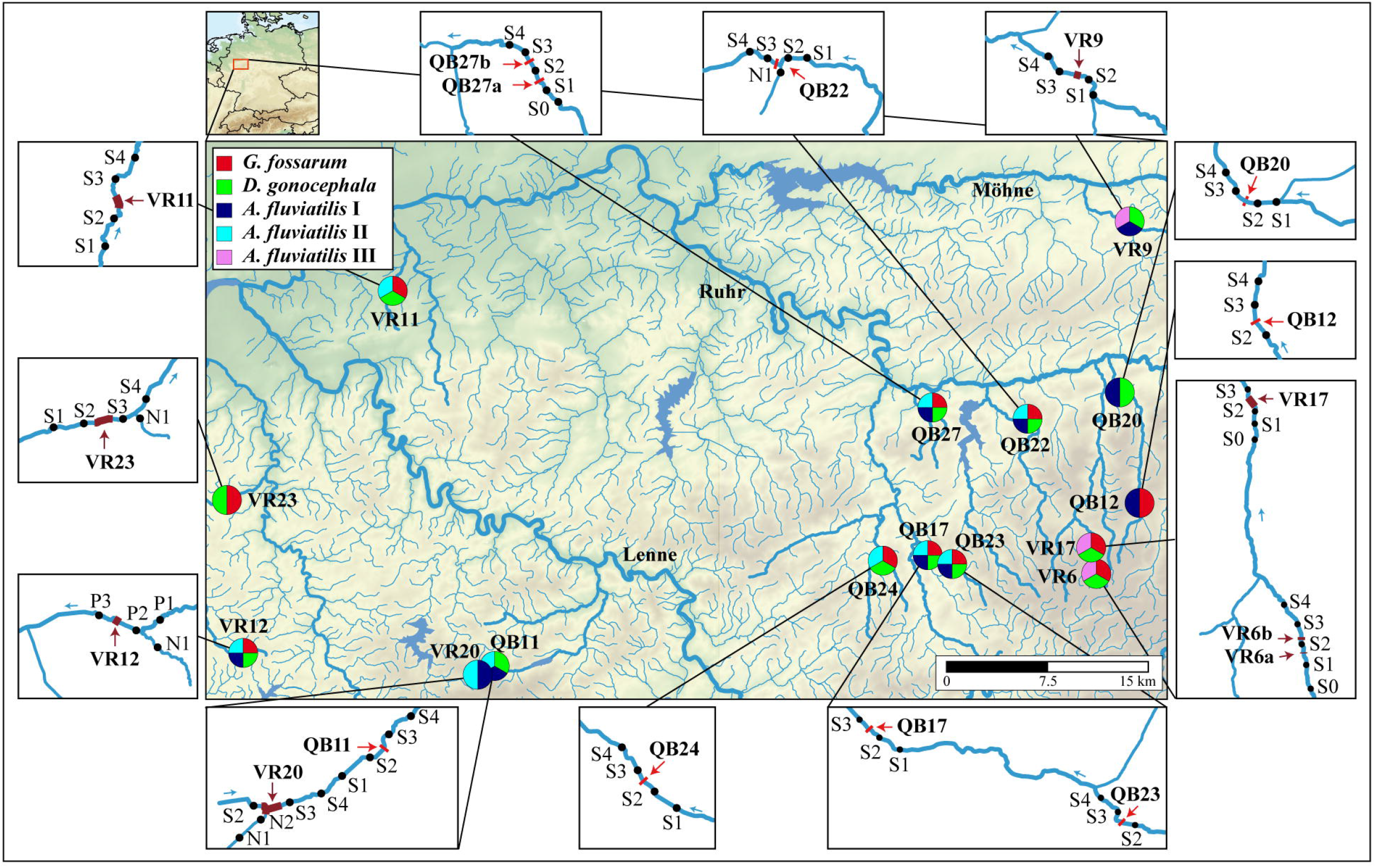
Central map showing the locations of barrier sites (QB = weirs, VR = culverts). Pie charts indicate sampled species. Water course of sampled streams and main rivers are highlighted. Boxes show the sampling scheme and locations of individual sampling sites, with a distance of approx. 200 m. Water flow direction is given by small arrows.

**Figure 3:**
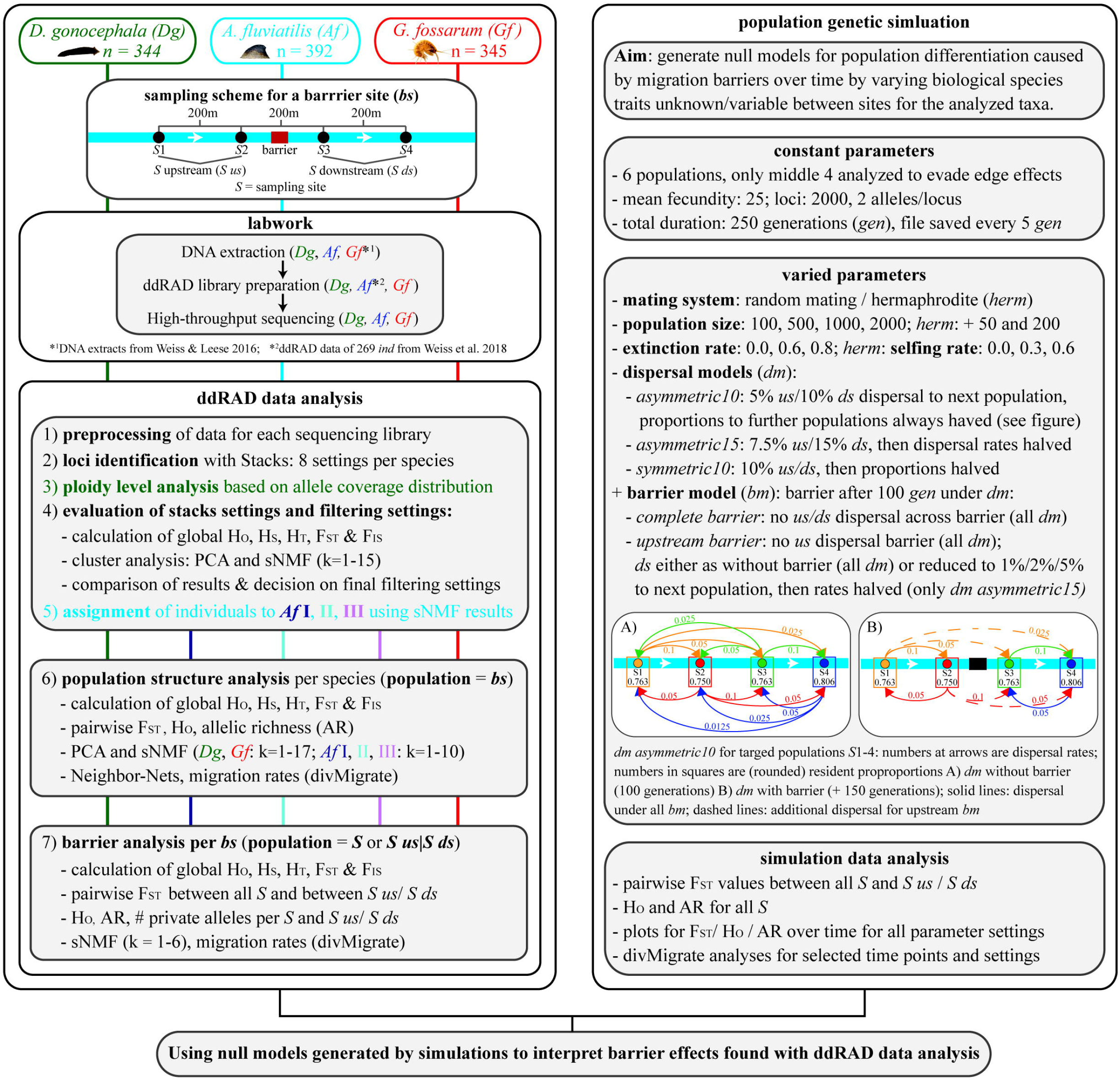
Workflow of the study. Left panel: Number of specimens analysed per species, sampling scheme for a barrier site, summary of lab work, bioinformatic and statistical analyses. For general population structure analysis per species (step 6), all specimens sampled at individual sampling sites before and after the barrier were pooled (barrier site). Right panel: Details on population genetic simulation including all important parameters and parameter space chosen for varied parameters and data analysis steps.

### Genotyping

DNA was extracted from muscle tissue using a salt-extraction protocol as described in Weiss & Leese (2016) for 344 *D. gonocephala* and 123 *A. fluviatilis* specimens. For *G. fossarum*, DNA from 315 specimens from a previous study (Weiss & Leese, 2016) was used. Four ddRAD libraries were generated for *D. gonocephala* (containing 83, 83, 87 and 89 samples), four for *G. fossarum* (92, 66, 65 and 95 (3 repeated from library 1)) and two for *A. fluviatilis* (62 and 61). For *A. fluviatilis*, data from three previously generated ddRAD libraries (269 specimens, Weiss et al., 2018) were also used, resulting in 389 specimens. The ddRAD libraries were generated according to the protocol described in Vendrami et al. (2017), with modifications described in Weiss et al. (2018) and Weigand et al. (2018). An overview of the protocol with species-specific details is given in Appendix S1 and sample preparation details are given in Table S2. All samples for each library were pooled with equimolar concentrations and sequenced on an Illumina HiSeq 2500 sequencer to obtain 125 bp paired-end reads (Eurofins Genomics Europe Sequencing GmbH; Constance, Germany).

### ddRAD data analysis

For a better overview, all important analysis steps are summarised in the workflow in Figure 3. Pre-processing of ddRAD data for each sequencing library was conducted as described in Weiss et al. (2018). After pre-processing, denovo_map.pl of Stacks v.1.34 (Catchen et al., 2013) was used to identify loci and genotypes. Stacks was run with eight different settings per species to identify the optimal parameter settings (Paris, Stevens, & Catchen, 2017, see Appendix S1 for details). Stacks results were exported from stacks databases with export_sql.pl (minimum stack depth 8). For *D. gonocephala,* aneuploid cytotypes are known besides diploids, with chromosome numbers similar to triploids (de Vries, 1986). Therefore, individual ploidy levels based on the expected allelic coverage was estimated using the R-script ploidyCounter.R (Rozenberg, https://github.com/evoeco/radtools/, see Weigand et al. 2018 for details). However, this ploidy estimation is indirect and aneuploids with high chromosome numbers cannot be distinguished from real triploids. Therefore, specimens with coverage distribution plots as expected for triploids are called ‘potential triploids’ hereafter. Direct estimation of ploidy levels, e.g. via flow cytometry was not possible from the ethanol fixed material. All analyses were also conducted excluding potential triploids, but as they produced similar results only analyses including all specimens are presented here.

Further analyses were performed using Snakemake workflows (Köster & Rahmann, 2012), combining stacks2fasta.pl (Macher et al., 2015) and several in-house python and R-Scripts for data reformatting, filtering and population genetic analyses. Parameter settings were similar for each species, except for *Ancylus* species, where parameters were adjusted when possible to account for polyploid genomes.

First, Stacks parameter and locus filtering settings were optimised, as described in detail in Appendix S1. General population structure was analysed with individuals divided into populations according to barrier sites, i.e. pooling specimens from individual sampling sites at a barrier site to one population (Figure 3, step 6). The following locus filtering settings were chosen: 1–12 SNPs/locus (only one used), minor allele frequency 1%, locus present in at least 80% of individuals per population and 95% of all individuals. Further, specimens with too few reads, or more than 20% missing data were excluded. Basic population genetic statistics, e.g. observed heterozygosity (*H*_O_), observed gene diversity (*H*_S_), overall gene diversity (*H*_T_) and overall *F*_ST_ and *F*_IS_ were calculated using the R-package hierfstat (Goudet, 2005) in R v. 3.3.2 (R Core Team, 2015). Further, principal component analyses (PCAs; Patterson et al., 2006) were performed and individual ancestry coefficients were estimated based on sparse nonnegative matrix factorisation algorithms (sNMF; Frichot et al., 2014) using the R-package LEA (Frichot & François, 2015) to identify genetic clusters. Both methods are suitable for polyploid and mixed ploidy data as they do not assume Hardy–Weinberg equilibrium (Dufresne et al., 2014; Frichot et al., 2014). For sNMF analyses, number of clusters tested were k = 1–15 for *Ancylus* species and k = 1–17 for *G. fossarum* and *D. gonocephala*, and analyses were run with 30 replicates and 100,000 iterations per replicate. In addition, Neighbour-Net networks (Bryant & Moulton, 2004) were calculated using SplitsTree v. 4.14.5 (Huson & Bryant, 2006). Pairwise *F*_ST_ values (after Weir & Cockerham, 1984) between barrier sites were calculated and significance was tested by bootstrapping over loci (10,000 replicates; 0.025/0.975 confidence intervals) using the R-package hierfstat. *H*_O_ and allelic richness (AR) in each population were calculated using the R-package diveRsity (Keenan et al., 2013). Further, the divMigrate function (Sundqvist et al., 2016) of this R-package was used to assess directional relative migration rates and to detect asymmetries in gene flow using Nei’s *G*_ST_ (Nei, 1973). To evaluate correlation of genetic and geographic distances (isolation by distance, IBD), Mantel tests were conducted using the R-package vegan (Oksanen et al., 2019). As a measure of genetic distance, pairwise *F*_ST_ values were calculated between all individual sampling sites (S). For geographic distances, either straight-line or waterway distances were used. Both distances were calculated using QGIS v. 2.14.14 (http://qgis.org) with the same stream map used for the visualisation of sampling sites and population structure, provided by the federal state authority LANUV (Gewässerstationierungskarte des Landes NRW © LANUV NRW (2013)).

Finally, the effects of different in-stream barriers on gene flow among populations up- and downstream of barriers were analysed (Figure 3, step 7). To enable detection of low differentiation among adjacent sites, loci were exported separately for each barrier site with the same filter settings used for previous analyses, except the minor allele frequency was set to 5% and loci had to be present in 90% of specimens to account for the smaller sample sizes. For barrier analyses, two population definitions were used. First, all analyses were conducted with barrier sites as individual populations, i.e. specimens from individual sampling sites up- and downstream of a barrier were pooled (S upstream/S downstream). Second, all individual sampling sites (S) were treated as different populations (see Figure 3). Basic population genetic statistics, sNMF (k = 1-6), *H*_O_ and AR per population and pairwise *F*_ST_ were calculated as before. The number of private alleles in each population was calculated using an in-house python script. To account for differences in sample size, larger populations were randomly subsampled 30 times to match the smallest population size and the mean number of private alleles among replicates was calculated. DivMigrate was used to calculate migration rates between sampling sites and to assess asymmetric migration patterns. A large number of loci (>1000) can partly compensate for the low sample sizes (Willing et al., 2012), but results for populations with n < 5 should be interpreted with caution (Sundqvist et al., 2016). To test the hypothesis that small in-steam barriers generally increase fragmentation compared to reference sites and that weirs have a stronger effect than pipes or tunnels (both culverts), linear mixed-effect models (LMMs) were run using the R-packages lme4 (Bates et al., 2015) and lmerTest (Kuznetsova et al., 2017). First, mean differentiation (*F*_ST_) for reference sites and across barriers were calculated per barrier and species. Different LMMs were run for each species with differentiation (*F*_ST_) as dependent variable, presence of barriers between compared sites as fixed effect and barrier site as random effect. In a second LMM analysis, barriers were subdivided into barrier types, i.e. weirs, pipes, and tunnels (and no barrier) and compared per species.

### Population genetic simulations

To simulate expected population differentiation caused by migration barriers over time, Nemo 2.3.51 (Guillaume & Rougemont, 2006) was used. As there is only little information available about the parameters needed to be given in the simulation, various barrier strengths and biological species traits were applied, including different mating systems (random mating for *G. fossarum* and hermaphrodite for *A. fluviatilis* and *D. gonocephala*), population sizes, extinction rates, selfing rates for hermaphrodites and dispersal and barrier models (see Figure 3 and Appendix S2 for details on parameter space). The dispersal and barrier models were designed to reflect stream dispersal (linear, influenced by flow direction) and our sampling scheme (four equidistant sampling sites). To prevent edge effects (no further dispersal possible for edge populations in the simulation), we simulated six populations but only analysed the four target populations. Simulations were run for 250 generations. The first 100 generations were simulated with different dispersal models but without a barrier to separate dispersal model and barrier effects. After 100 generations, a barrier was introduced for the next 150 generations using three models (complete barrier, upstream barrier with unchanged, or reduced downstream dispersal, see Figure 3 for details) and fstat files were saved every five generations.

Data analyses were conducted using different python and R-Scripts in a Snakemake workflow. Edge populations were excluded and ten individuals per population were randomly chosen to reflect average number of specimens sampled per population. Populations with fewer than ten individuals at specific time points were excluded. Pairwise *F*_ST_, *H*_O_ and AR were calculated for all populations every five generations for each parameter combination, as described for ddRAD data. *F*_ST_ values were calculated for all single populations and for pooled populations on both sides of the barrier. To evaluate the performance of divMigrate in the detection of reductions of migration rates due to barriers, a subset of output files were generated with the migration model “*asymmetric15*” and all barrier models (except upstream barrier with reduced downstream 5%) for both mating systems and population sizes of 100 (500 for reduced downstream dispersal models) and 1000 with extinction rates of 0 or 0.6. Migration rates (using *G*_ST_) were calculated at different time points before (5, 50 and 95 generations) and after barrier introduction (after 110, 150, 200 and 250 generations).

## Results

All five macroinvertebrate species showed strong regional and local population structure as detected by all methods.

### Population structure of *Gammarus fossarum*

We found *G. fossarum* in nine streams at 11 barrier sites (Table S1, Tables S3 and S4 for summary statistics for all species). PCA (Figure S2) and sNMF analyses indicated strong population structure. In the sNMF analysis, cross-entropy (CE) was lowest for k = 9 (Figure S3A). The nine clusters corresponded to the nine streams, also visible in the Neighbour-Net analysis (Figure 4A and B). Further, the network showed superordinate clustering into three phylogenetically distinct groups in concordance with sampling site topography. Pairwise *F*_ST_ values were generally high (mean 0.47), especially between network-groups and all values were significant, including values for the comparison between barrier sites in one stream, ranging between 0.01 and 0.74 (Figure 4C). Likewise, estimated migration rates were very low in most cases (mean 0.05), with the exception of within-stream comparisons (Figure 4D, Table S5A). However, for the two within-stream comparisons, migration rates between VR6 and VR17 were much lower in both directions than those between QB17 and QB23 and were significantly higher in the downstream than in the upstream direction. In 71% of the comparisons, significant asymmetric gene flow was detected.

**Figure 4.**
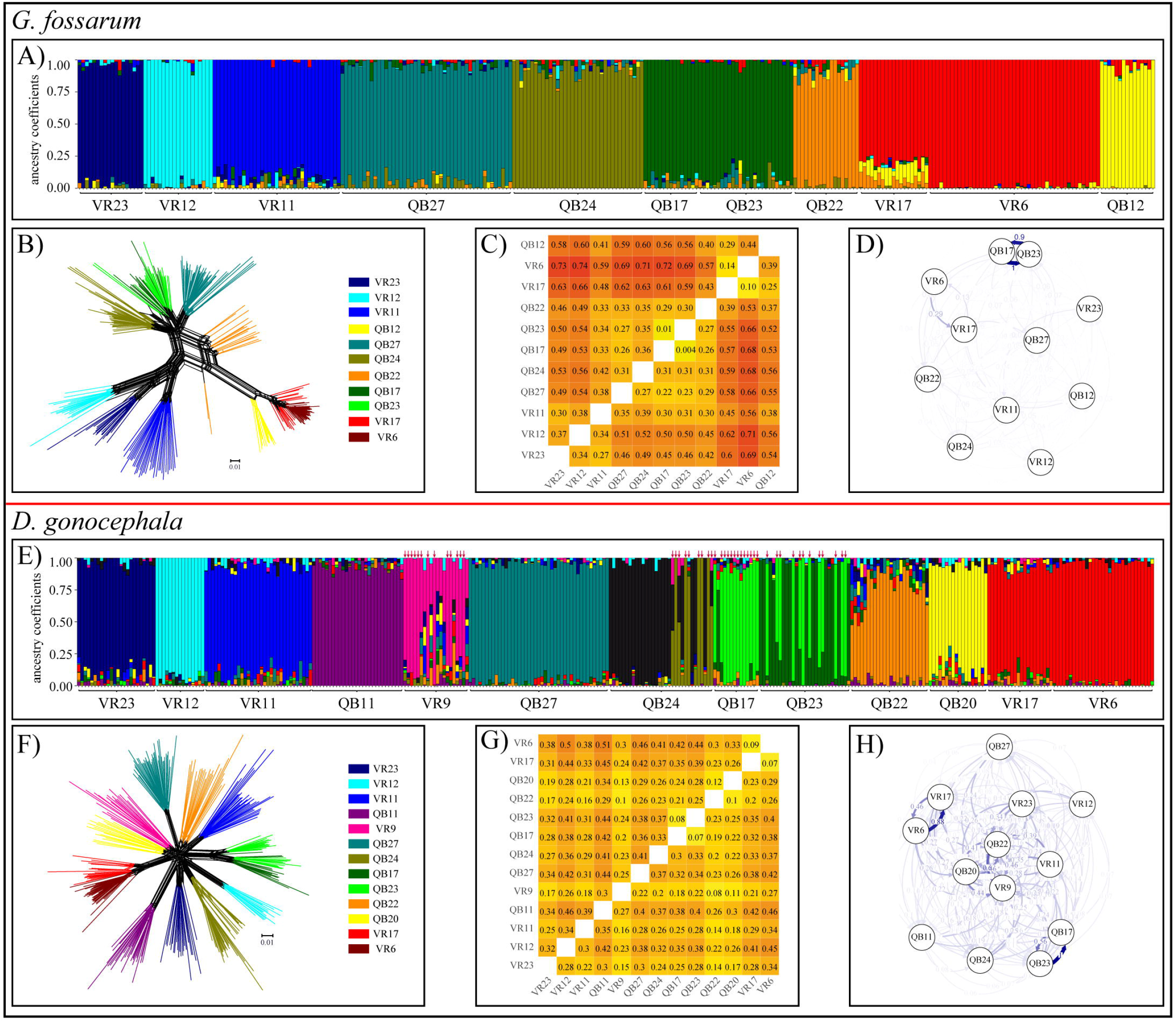
Population genetic structure of *G. fossarum* (A-D) and *D. gonocephala* (E-H). A, E) Ancestry estimates for A) K = 9 and E) K = 13, where vertical bars represent individual ancestry coefficients. B, F) Neighbour-Net, branch colours indicate barrier sites. C, G) *F*_ST_ heat maps for pairwise comparisons between barrier sites. Above the diagonal pairwise *F*_ST_ values are given and below the lower confidence interval (values >0 indicate significant differentiation), all coloured according to the level of differentiation. D, H) DivMigrate plot, strength and visibility of arrows are correlated with migration rates.

### Population structure of *Dugesia gonocephala*

*D. gonocephala* was found at 13 barrier sites in 11 streams. Of 344 specimens, 294 were diploid and 50 were potentially triploid. PCA (Figure S2) and sNMF cluster analysis both indicated strong population sub-structuring. CE was lowest for K = 13 (Figure S3B). The population structure could be explained best by the stream origin and ploidy level (Figure 4E). At sites with both diploids and potential triploids (QB24, QB17 and QB23 and VR9), individuals were divided into two clusters or showed high intermixture between clusters. The Neighbour-Net showed a star-like pattern containing 11 clusters, each representing a single stream (Figure 4F), but sub-clusters within streams according to ploidy level were visible. *F*_ST_ values were lower than those for *G. fossarum* (mean: 0.31, range 0.08 to 0.51) but significant for all pairwise comparisons including within stream comparisons (Figure 4G). In general, estimated migration rates were higher than those for *G. fossarum* (mean: 0.16) and migration rates for most comparisons were significantly asymmetric (73%; Figure 4H, Table S5B). Similar to *G. fossarum*, migration rates were highest between sites within a stream and migration was significantly higher in the downstream direction.

### Population structure of *Ancylus fluviatilis* I, II and III

Among 389 specimens, 167 were assigned to *A. fluviatilis* I, 159 to *A. fluviatilis* II and 63 to *A. fluviatilis* III. In contrast to *G. fossarum* and *D. gonocephala*, *H*_O_, *F*_ST_ and *F*_IS_ values varied strongly between stacks settings (Table S4). For *A. fluviatilis* I, PCA (Figure S2) and sNMF indicated strong genetic structure. CE values were low k = 7–10 (Figure S3C), with the lowest mean CE over repetitions for k = 8. The clusters mainly corresponded to streams, but more specimens showed higher admixture to additional clusters (Figure 5A). These results corresponded well with the network results (Figure 5B), where a similar group structure consistent with streams was visible, with the exception of specimens from QB12 and QB20, which were split into two groups according to the barrier site. In general, pairwise *F*_ST_ values were lower than those in *G. fossarum* and *D. gonocephala* (mean: 0.18, range: 0 – 0.29, Figure 5C), but all were significant with the exception of the within-stream comparison QB11 and VR20. Migration rates were between the two previously described species (mean: 0.11), with 80% of comparisons showing significant asymmetry (Figure 5D, Table S5C).

**Figure 5.**
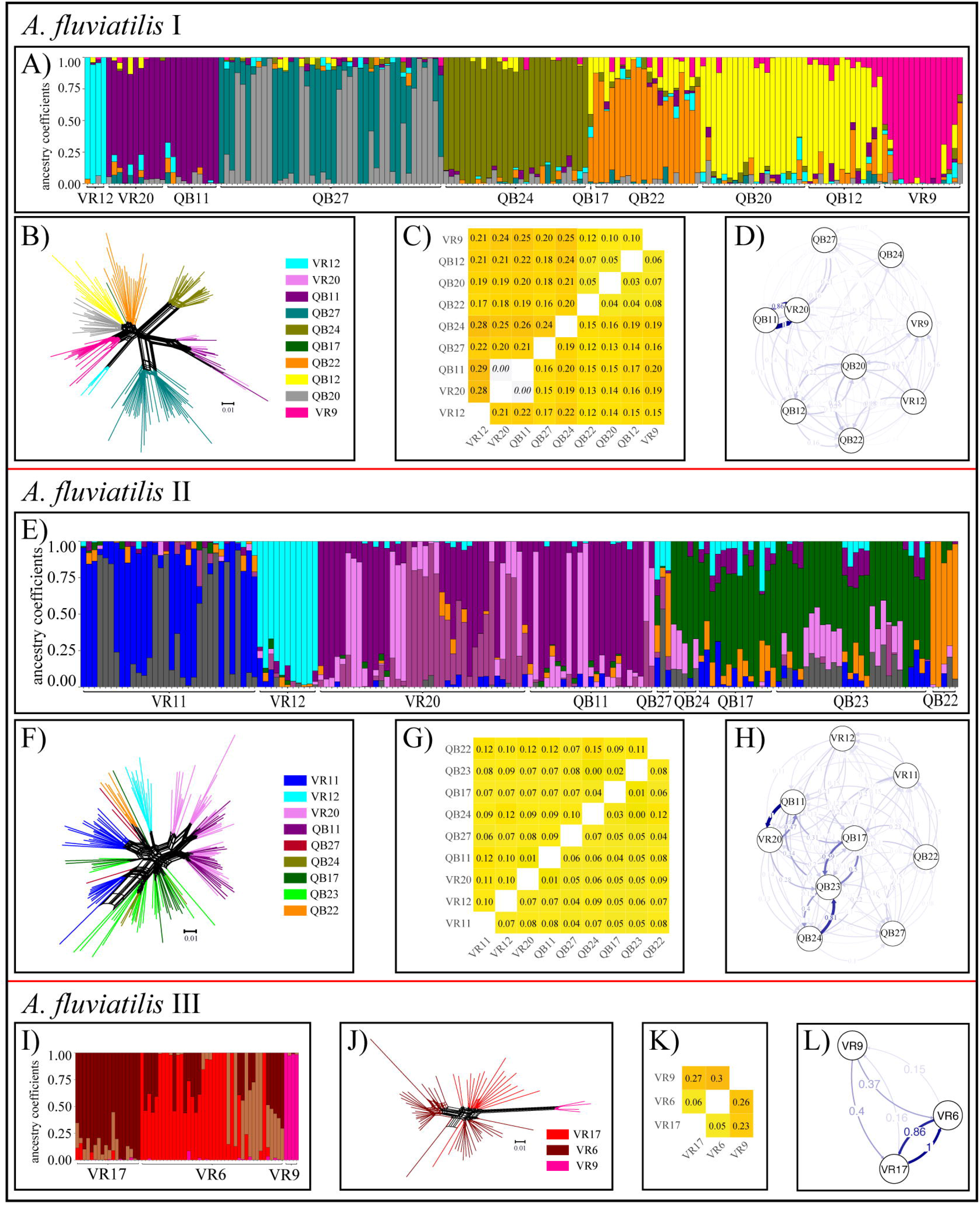
Population genetic structure of *A. fluviatilis* I (A-D), II (E-H) and III (I-L). A, E, I) Ancestry estimates for A, E) K = 8, I) K= 4, where vertical bars represent individual ancestry coefficients. B, F, J) Neighbour-Net, branch colours indicate barrier sites. C, G, K) *F*_ST_ heat maps for pairwise comparisons between barrier sites. Above the diagonal pairwise *F*_ST_ values are given and below the lower confidence interval (values >0 indicate significant differentiation), all coloured according to the level of differentiation. Non-significant values and corresponding CI are indicated in italics. D, H, L) DivMigrate plot, strength and visibility of arrows are correlated with migration rates.

For *A. fluviatilis* II, PCA (Figure S2) and sNMF both indicated strong population structure within the species. The most probable number of clusters in the sNMF analysis was k = 8 (Figure S3D). The clusters were less clear than those for the other species, as many specimens showed high amounts of shared ancestry to different clusters. The main factor explaining the groups was stream origin (Figure 5E), but in some cases, main cluster were shared among different streams or additional cluster were found within one stream. A similar structure was observed in the Neighbour-Net analysis. However, single specimens with high admixture in the sNMF plot, grouped together with specimens sampled in other streams (Figure 5F). Overall, pairwise *F*_ST_ values indicated relatively low but significant differentiation between all sampling sites, except for QB23–QB24 (mean: 0.08, range: 0.003 – 0.15, Figure 5G). Estimated migration rates were higher than those for the other species (mean: 0.17) and less asymmetry was detected (58% of comparisons, Table S5D).

*A. fluviatilis* III was mainly found in one stream with two barrier sites (VR6 and VR17). At VR9, four individuals were identified as *A. fluviatilis* III/I hybrids (75%/25%) according to Weiss et al. (2018) and these were included here. In the PCA (Figure S2) and sNMF analysis, sub-structure was detected, with four clusters best explaining the population structure (Figure S3E). One cluster consisted of VR9, the second of VR17 specimens and the other two of VR6 specimens, with shared proportions for some individuals between clusters (Figure 5I). A similar pattern was observed in the Neighbour-Net analysis (Figure 5J). Pairwise *F*_ST_ values were all significant but lower between VR6 and VR17 (Figure 5K), where also higher migration rates were detected (Figure 5L, Table S5E).

For all species, we detected a correlation between geographic and genetic distances, which was strongest for *A. fluviatilis* II and weakest for *D. gonocephala* (Figure S4). Genetic diversity (i.e., *H*_O_ and AR) differed among sites for all species, but with no consistent pattern over all species (Table S6).

### Effects of in-stream barriers on population structure

Filtering loci separately for each barriers site, resulted in in a greater number of variable loci and mostly increased genetic diversity per barrier site in comparison to the total dataset (Table S7). We could not detect distinct barrier effects across species when comparing *F*_ST_ values between reference and barrier separated sites. Mean differentiation was low and mostly insignificant regardless of the presence of barriers or the barrier type (weir, pipe or tunnel). Consequently, none of the LMMs indicated a general effect of barriers on differentiation (Appendix S3). When studying specific barriers in detail, we found no evidence for a barrier effect on population structure for seven of the nine weirs and six of the eight culverts in any species. This means that populations at barrier sites were either i) not differentiated at all, ii) differentiation was similar among reference and barrier separated sites, iii) lower for some of the comparisons of barrier separated sites compared to reference sites, or iv) that only individual sampling sites were constantly differentiated to the other sites from the same barrier site. All these four differentiation patterns were found for all species. Differentiation between individual sites was most prominent for specimens sampled in affluents (“N”; Figure 6), despite the lack of obvious barriers. At three weirs, reduced upstream migration rates were detected for *A. fluviatilis* I and II (QB11, QB12 and QB17) (Figure S5). However, we found no indication for a barrier effect in any of the other analyses. A similar reduction was observed for VR6a for *D. gonocephala* but here the sample size was too small at upstream sites (Figure S5).

**Figure 6.**
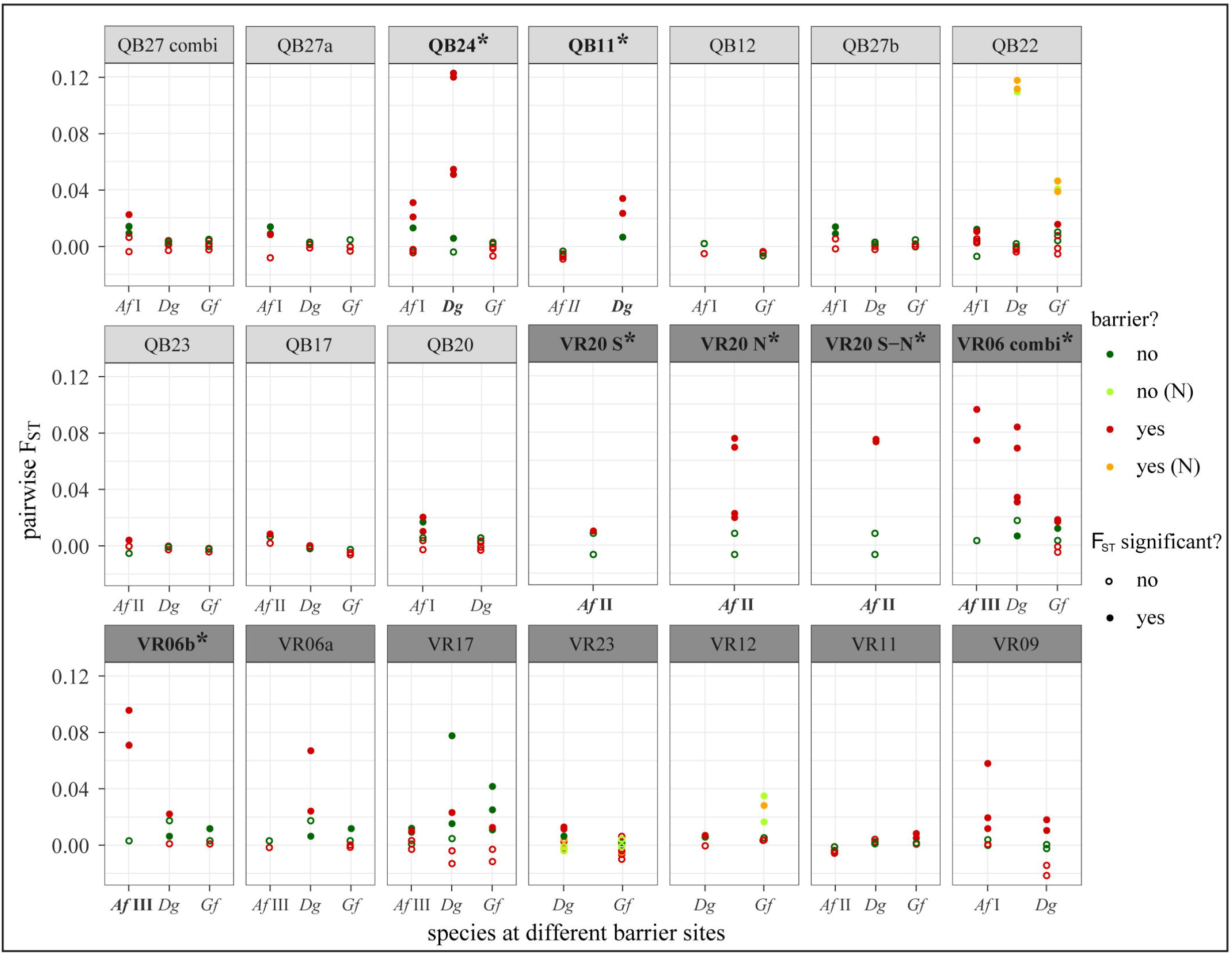
Pairwise *F*_ST_ values between all sampling sites for each barrier and species, coloured according to presence of barriers. *F*_ST_ values from site stream populations (N) are shown in different colours. Barriers are ordered according to severity, starting with the strongest weir and strongest culvert. Combi: differentiation across both barriers at one barrier site. Barriers where an effect was detected are indicated with an asterisk.

We only detected distinct barrier effects associated with weirs for *D. gonocephala* at QB24 and QB11. In both cases, differentiation across barriers was larger than that between reference sites (Figure 6). The effect was stronger for QB24 than for QB11, consistent with the sNMF results, where two clusters were the most probable for QB24. Specimens sampled upstream of the barrier only belonged to the first cluster, while specimens assigned to both clusters, as well as intermixed ones, were found downstream of the barrier. Specimens showing high membership to the second cluster were all classified as potential triploids. At both barriers, migration rates in up- and downstream directions were lower across barriers than among reference sites. For QB24, all rates across the barrier were significantly asymmetric, with stronger downstream dispersal. Also, genetic diversity was higher downstream of barriers at both sites, indicated also by higher amounts of private alleles (Tables S8 and S9).

A significant barrier effect was detected for *A. fluviatilis* II at VR20 and for *A. fluviatilis* III at VR6b. While VR6b was a pipe, VR20 was a combination of a pipe and tunnel with an additional steep drop within and two streams were joined within the tunnel. At VR20 S4, we only found one specimen belonging to *A. fluviatilis* II, which was excluded from analyses of *F*_ST_ and migration rate. The other 11 specimens at this site were identified as *A. fluviatilis* I, which was not detected upstream of the culvert. All individual site comparisons across the culvert in *A. fluviatilis* II indicated significant differentiation, including comparisons of both N1 and N2 with S2, which were separated by a pipe and part of the tunnel, while differentiation among reference sites was not significant. This pattern was also detected in the divMigrate analysis. Upstream migration across the culvert to both ends was highly reduced, while downstream migration rates were similar or even higher than those among reference sites (Figure S5). In the sNMF analysis, indications for a barrier effect were found with the lowest CE for three clusters. Specimens from both N1 and N2 were split into two clusters (k1: n = 10, k2: n = 6), S1 specimens were assigned to k2 or the third cluster (k2 = 6, k3=5) and specimens at S3 were assigned to all three clusters, indicating that downstream dispersal was possible. AR was lowest at S2; otherwise, genetic diversity was similar (Tables S8 and S9).

At barrier site VR6, the longer pipe of this barrier site (VR6b) caused population differentiation in *A. fluviatilis* III, as all comparisons across the pipe were significant and *F*_ST_ values were higher than those between reference sites (Figure 6). These differentiation patterns were supported by the sNMF analysis, where three clusters were detected. One cluster was only found in S3&4, while the other two were mainly found in S1&2 and less frequently in S3&4, where assignment to multiple clusters was observed. While migration rates between sites were generally reduced, upstream migration across the second pipe (VR6b) was even stronger reduced to nearly zero. By contrast, migration rates across the first pipe (VR6a) were even higher than those among reference sites (Figure S5). *H*_O_ was similar at all sampling sites, while AR and the number of private alleles were higher at S3&4 than at other sites (Tables S8 and S9).

Unlike in *D. gonocephala* and *A. fluviatilis*, we did not detect evidence for a barrier effect on population structure in *G. fossarum* at any of the barriers.

### Population genetic simulation

The simulation of a complete barrier revealed the quick emergence of strong population differentiation, as determined by increases in *F*_ST_ values and decreases in migration rates over time (Figure S6, Table S10). In general, barrier effects were similar, independent of the distance between populations. In smaller populations, fluctuations over time were larger, leading to low but significant population differentiation between reference populations not separated by a barrier. However, after only a few generations, *F*_ST_ values for populations separated by a barrier were consistently higher than those between reference populations. For larger populations (i.e. patch capacity 2000), it took up to 50 generations to detect differentiation and *F*_ST_ values remained low but followed the same general pattern (Figure S6). Here, only *F*_ST_ values between populations separated by a barrier were significant. The higher the extinction rate, the higher the *F*_ST_ values between barrier-separated populations and also in larger populations, differentiation was detected earlier. For smaller populations with higher extinction rates, significant differentiation was detected without a barrier but remained lower than for barrier-separated populations. These patterns were visible for all tested migration matrices, with asymmetric matrices leading, on average, to slightly higher *F*_ST_ values. However, while a reduction of migration rates over time was detected with the divMigrate approach, the simulated asymmetry was not reflected in consistent differences between up- and downstream migration rates. Applying different selfing rates to simulate hermaphrodites, *F*_ST_ values increased with increasing selfing rates, but patterns were generally the same as those for populations with obligate sexual reproduction. Even though complete isolation between populations on both sides of the barrier was not directly detected, a reduction in migration rates after barrier introduction was quickly (<10 generations) detected for all migration scenarios and decreased further with time. The reduction was more easily detected for larger sample sizes, higher extinction rates and smaller populations.

In general, barriers impeding upstream but not downstream dispersal did not affect the population structure and did not result in reduced inferred migration rates (Figure S6, Table S10). At some time points for small populations (particularly in combination with higher extinction and/or selfing rates), *F*_ST_ values were slightly higher and significant for barrier-separated populations than for reference populations, but this pattern fluctuated over generations and did not increase with time (similar for migration rate estimates). Reducing downstream dispersal rates led to an intermediate pattern between those described previously. Not all parameter combinations were simulated, but downstream dispersal rates had to be reduced substantially to 1 %, or depending on the other parameters 2 %, to influence population differentiation. The effects were stronger (and sometimes only detectable) for smaller population sizes, higher extinction or selfing rates, and were strongest for a combination of these parameters (i.e., if population sizes were small and extinction rates were high). If the selfing rate was increased from 0.3 to 0.6, the effect on the population structure was generally high but barrier effects were less detectable because fluctuations between populations and over generations were higher. We obtained similar results for migration rates, but correct asymmetry was not reliably detected. *H*_O_ and AR were, in general, not affected by the barrier model (Figure S7). Both AR and *H*_O_ decreased stronger over time for smaller population sizes and higher extinction proportions for all dispersal models (with no or slow decreases for larger population sizes, i.e., 1000 and 2000). However, the decrease was constant over time, independent of the introduction of a barrier into the system and similar in all populations.

## Discussion

### General effects of small in-stream barriers on macroinvertebrate taxa

For all five stream species, our population genomic data revealed strong population differentiation at local scales. However, in contrast to our hypothesis, the tested individual small barriers did only increase genetic differentiation compared to control sites in few cases. No correlation between barrier strength (weir, pipe or tunnel) and differentiation was found. Therefore, we found little evidence that the tested in-stream barriers are the cause of the observed local scale population isolation. Only for few individual in-stream barriers we detected a significant barrier effect: two weirs for *D. gonocephala* and one culvert each for *A. fluviatilis* II and III. This raises the questions i) whether barrier effects are detectable with our chosen methods for barriers with an age of 33-109 years and ii) whether population genetic signatures indicate fragmentation if only upstream dispersal is impeded. To address both questions, we simulated population genetic null models under specific life history traits and explored life-cycle and barrier parameter space as realistic as possible. However, greater knowledge on the biology of the species would be needed to obtain more precise null models for comparison as otherwise endless possibilities for parameter combinations exist.

The simulation results consistently showed that barriers preventing up- as well as downstream dispersal lead to strong population structure relatively quickly within the age range of the barriers tested. Other studies simulating the effect of barriers on population structure found that barrier effects will only be detectable with *F*_ST_ after longer times (Landguth et al., 2010), or for small populations (Ne < 100; Coleman et al., 2018), but simulation parameters were quite different in these simulations, for example different sampling schemes or marker were used, making it difficult to compare simulations. Here, we found that the time until effects were detectable depended on the population size and extinction rates, with smaller populations showing barrier-related structure already after 5 to 20 generations and the largest populations (patch capacity 2000) after approximately 50 generations.

Our simulations indicated that *F*_ST_ was the most reliable indicator of fragmentation. *H*_O_ or AR could not be used as both decreased constantly over time in small reference and barrier-separated populations, or remained relatively constant in larger populations. With migration rate estimates by divMigrate, complete barriers were detectable as well by a general reduction in migration rates between barrier-separated sites. However, in general, migration rates were overestimated, suggesting that inferred low migration rates in real data probably indicate real barrier effects.

To conclude, simulations showed that even minor migration (2 % or 5 % depending on population size, extinction and selfing rate) in one direction can be sufficient to counteract barrier-related population differentiation. If populations are large (patch capacity >2000) and no extinction rates are simulated, even 1 % downstream migration can be enough to counteract differentiation in random mating populations. Accordingly, reduced migration due to barriers is expected to be detectable if migration rates are low, but detection will be difficult or impossible for very young (<10 years) barriers, for barriers only reducing upstream dispersal and for large effective population sizes.

### Species-specific barrier effects

We hypothesised that effects would depend on barrier strength and would be strongest for *A. fluviatilis*, intermediate for *D. gonocephala* and lowest for *G. fossarum*. Our results only partly support the hypotheses as only single species at a few strong barriers we detected a stronger population structure and reduced migration rates compared to those for reference populations. For most weirs and culverts, no barrier effect was found. This indicates that all species can in principle overcome the tested small in-stream barriers frequently enough (or at least in one direction) to facilitate gene flow. However, it cannot be ruled out that barrier effects were not detectable even though they exist because of too large effective population sizes or because barriers only reduced upstream dispersal as discussed above. Additionally to active dispersal, hololimnic specimens may overcome barriers by passive dispersal via animal vectors such as waterfowl. This process can be frequent at local scale (Coughlan et al., 2017) and experimental evidence suggests that it could be an important dispersal mechanism for amphipods, snails and other invertebrates (Rachalewski et al., 2013; van Leeuwen & van der Velde, 2012; Waterkeyn et al., 2010). While no information is available on the probability or frequency of zoochorous dispersal for the taxa studied here and in general more information is needed, e.g. on the quantitative contribution of zoochory to dispersal (Coughlan et al., 2017), it should be considered as a possible mechanism counteracting barrier effects.

With respect to the individual species, the barrier effects were weakest for *G. fossarum* clade 11, as expected. This indicates that dispersal ability is high enough to overcome the small in-stream barriers analysed here (see also Weiss & Leese, 2016), congruent with findings for *G. fossarum* clade 12 (Alp et al., 2012). It is also possible that the tested barriers were not old enough to create detectable isolation patterns (Monaghan et al., 2001), that the reduction in migration was not strong enough to impact population structure determined by the genetic markers (Whiterod et al., 2017), or that effective population sizes were too large as *G, fossarum* is expected to have the largest N_e_ of all taxa tested here. However, based on the barrier ages (>30 years, ≥ 30 generations), distinct local population structure, simulation results and the comparably high mobility of the species with the capability for active and passive dispersal, we consider it more likely that the tested small in-stream barriers do not present severe dispersal barriers for this species.

Our prediction that barrier effects would be moderate for *D. gonocephala* and strongest for *A. fluviatilis* was not supported by the data. Only two weirs and two culverts had species-specific barrier effects. The weirs influencing *D. gonocephala* were classified as the second (QB24) and third (QB11) most severe according to height, steepness and smoothness. They were both >80 years old. For other barriers of the same age, no population structure was detected as well as for QB27, which was even higher than QB24 with a smoother ramp slope. The difference between barriers of the same age might be explained by differences in population size or by important barrier characteristics that had not been measured. The detected barrier effect was weaker for QB11 than for QB24 which was not as high and had a moderate slope with drops at the start and the end. The two drops might have strengthened the effect in comparison to other barriers with similar slope. In general, our results indicate that *D. gonocephala* maintains gene flow across most small weirs, but we also found evidence that weirs of the size and shape tested can impede migration. This holds true especially for those classified as more or most severe. Thus, larger barriers, especially larger drops, will probably impact on dispersal for this flatworm species. Contrary to our expectations, *A. fluviatilis* migration was not stronger affected by weirs. At three weirs, upstream migration rates across barriers were lower than those between reference sites, but no effect was detectable with any other method. To finally conclude that small weirs do not influence migration in *A. fluviatilis*, it would be important to increase the sample size and to evaluate the influence of polyploidy on analyses. In addition, especially larger drops should be investigated as they will probably pose a greater barrier to dispersal than steep ramps, if they cannot be crossed by crawling or passive zoochorous dispersal.

For culverts, patterns were more consistent with our expectations. We only detected effects for *A. fluviatilis* II and III at two culverts predicted to have the strongest impact (VR20 and VR6b). At VR20, only *A. fluviatilis* II was found upstream of the culvert, while *A. fluviatilis* I was additionally found at downstream sites. While downstream dispersal seemed to be possible through this culvert, upstream dispersal was highly limited. A similar pattern was found for *A. fluviatilis* III at VR6b, even though here, inferred migration in both directions was reduced also between reference sites, but stronger across the pipe. For the other species, we could not attribute observed population structure between sampling sites to culverts, or the sample size upstream of the barrier was too small *(D. gonocephala*). These results indicate that all studied species can probably disperse effectively through tunnels of up to 120 m length, but pipes < 25 m can act as strong dispersal barriers probably due to their smooth internal structures and high flow velocity (David et al., 2014), or drops at the end, preventing upstream movement (Vaughan, 2002). Also for culverts it is possible that barrier effects went undetected due to, e.g. large N_e_. But in general our results were in accordance with the expectations that tunnels present less severe barriers than pipes, and it seems possible that the studied taxa can overcome them actively. However, with respect to pipes our results suggest that these can pose severe dispersal barriers even when relatively short. Therefore, it would be important to focus on this kind of culverts in further studies.

In general, single small barriers did not have a large impact on population structure, yet it remains to be determined whether cumulative effects exist and population structure increases when many small barriers occur within short distances. Future work should focus on testing and adapting novel and promising indices to identify population fragmentation in streams developed based on microsatellites (Prunier et al., 2020) to SNP-based data including also possible asymmetric gene flow and focus on barriers at the upper level of the severity gradient tested here, for weirs as well as for culverts.

### Regional subdivision

While effects of individual in-stream barriers was minor, genetic differentiation between streams was high for all species despite the small spatial scale of the study. Based on life history traits, we hypothesised that population structure to be weakest for *G. fossarum*, intermediate for *D. gonocephala*, and greatest for *A. fluviatilis*. However, contrary to our expectation, *G. fossarum* showed the highest population differentiation and the lowest migration rates between populations, *A. fluviatilis* species showed the lowest and *D. gonocephala* showed an intermediate level of differentiation.

For *G. fossarum*, populations were already highly differentiated at the shortest distances of 11– 14 km, with *F*_ST_ values of approximately 0.35. Analysing populations sampled at sites approximately 2 km apart within a single stream showed that gene flow in *G. fossarum* is possible over this distance but can already be reduced. In contrast to the other species, we also detected a strong phylogeographic structure. Populations clustered into three groups in the network analysis, congruent with the geographic distribution and mitochondrial COI groups defined by Weiss & Leese (2016), indicating that the area was probably recolonised by at least three distinct historical source populations after Pleistocene glaciation. This strong influence of historical processes on population differentiation could explain the higher population differentiation in comparison the other species. Still, the strong differentiation after 10 km is inconsistent with the relatively high mobility, locally often large populations and regionally broad distribution. It is possible that the dispersal ability is lower than expected or, more likely, that unidentified obstacles to dispersal exist. These obstacles could be human-induced alterations and fragmentations, such as larger dams, weirs and culverts (especially larger pipes), or unfavourable conditions in connecting areas, such as those caused by anthropogenic land use, organic pollution, acidification or large connecting rivers, or a cumulative effect of these factors (e.g., Alp et al., 2012; Cook et al., 2007; Monaghan et al., 2001; Watanabe et al., 2010). Furthermore, gene flow does not correspond directly with individual movement of specimens (e.g., Bohonak & Jenkins, 2003) when dispersing individuals are not able to successfully establish in an existing population. This phenomenon is described by the monopolisation hypothesis (e.g., De Meester et al., 2002), which predicts that adaptation along with the numerical advantage of first migrants leads to a priority effect of the founder population over new migrants and therefore reduces establishment success. Such intraspecific priority effects as well as isolation by adaptation patterns have been detected in various taxa (e.g., Boileau et al., 1992; Fraser et al., 2014; Nosil et al., 2008; Urban & De Meester, 2009) and could have led to increased differentiation.

In *D. gonocephala*, we detected fairly high population differentiation, but in contrast to *G. fossarum*, no strong geographic structure was found, supported by the low IBD pattern. Populations separated by only approximately 2 km in the same stream already showed significant population differentiation. Differentiation was also detected for 11 to 18 km but was not correlated with waterway distance within this range, suggesting that additional factors other than waterway distance influence effective migration in this species. Local adaptation could shape the population structure, as reported for a population in the same study area in response to high copper concentrations (Weigand et al., 2018), in which also high population differentiation was found. Apart from this, little is known about effective dispersal in *D. gonocephala*, but planarians are generally been regarded as weak dispersers (e.g., Rader et al., 2017). The overall lower *F*_ST_ values in comparison to *G. fossarum* could indicate that *D. gonocephala* is a better disperser. However, it is more likely that differences in *F*_ST_ reflect the greater phylogenetic signal in *G. fossarum*, represented by three deep phylogenetic lineages as described above. In *D. gonocephala*, particularly strong population structure in some cases was associated with the presence of potentially triploid individuals. Potential triploids originating from different streams did not cluster together but were closely related to specimens in the same stream, indicating that polyploidy evolved independently several times. However, results concerning possible polyploid individuals have to be interpreted with caution, i) because it is difficult to assess how analyses are affected for example by allelic dosage uncertainty (Dufresne et al., 2014) and ii) because we estimated ploidy levels only indirectly and cannot say if individuals are really triploids or aneuploids with high chromosome numbers as it has been implied for a French *D. gonocephala* population (de Vries, 1986). Therefore, chromosome numbers in respective populations should be analysed by karyograms or flow cytometry. At sites with potential triploids, they often made up the majority, consistent with previous results showing that asexual polyploids are more abundant than diploids (Álvarez-Presas & Riutort, 2014). Asexual populations of *D. gonocephala* have not been reported (Stocchino & Manconi, 2013), but we cannot exclude the possibility and polyploid populations should be analysed in more detail in future studies.

The low genetic structure in *A. fluviatilis* species could be explained by the presence of three relatively young cryptic species in the area (see Weiss et al., 2018 for details), as time since speciation might have been not long enough to generate similar variation compared to e.g. the extremely diverse *G. fossarum*. Further, analyses were complicated by the polyploid genomes. The chosen analysis approaches should be able to generate reliable results concerning population genetic structure (Dufresne et al., 2014), but also loci detection could have been influenced by the ploidy. Even though we chose the most rigorous Stacks settings to disentangle homoeologous loci of different genomes (as defined in Dufresne et al., 2014), observed heterozygosity, particularly for *A. fluviatilis* II, was still much higher than for *D. gonocephala* and *G. fossarum*. Even though this filtering was necessary, it led to fewer loci being analysed for the different *A. fluviatilis* taxa. Despite the weaker population structure, significant differentiation between populations originating from different streams and generally low migration rates among populations were detected for all *A. fluviatilis* species and a high number of genetic clusters was supported by the sNMF analyses. Additionally, most within-stream comparisons showed low but significant differentiation, even within a few hundred meters. In general, our findings suggest that effective dispersal is low between streams and differentiation within streams can occur over short distances (see also Macher et al., 2016). In some cases, we detected genetic structure even within barrier sites, while in other cases, levels of shared ancestry between clusters were higher than those for *G. fossarum* or *D. gonocephala*. This may reflect higher occasional gene flow between streams, a younger recolonisation history, or could have been influenced by polyploidy and the still high heterozygosity in the data set. For a better understanding of the determinants of population structure, it is important to better characterise homoeologous loci and to determine ploidy levels of individuals. However, the consistent overall patterns across species suggest that our results are reliable and population isolation despite differences in dispersal capacity indicates that strong dispersal limitations exist in the area, which are not identified, yet.

## Conclusions

We detected strong genetic isolation among populations of five hololimnic species between streams. While for some individual barriers and species an effect was found, our genomic ddRAD data suggest that single small weirs and culverts are probably not the cause for the detected strong fragmentation at few kilometres distances. It remains to be tested, if cumulative effects of small barriers could have caused the disruption of gene flow between populations, or if other factors, such as larger in-steam barriers, land use, chemical pollution, urbanisation, or a combination of these, led to the observed population structure. In general, our combination of genomic markers, population genetic simulations and a controlled sampling design with distinct reference populations is a suitable tool to infer major drivers of regional and local population structure.

## Acknowledgements

We thank Lisa Pöttker, Markus Patschke, Vivienne Dobrzinski, Janis Neumann and Tobias Traub for assistance with sampling; Ralph Tollrian for support; Romana Salis for helpful comments on the manuscript. Furthermore, we thank Jörg Drewenskus (Bezirksregierung Arnsberg) for information on barrier ages and LANUV for further information on sampling sites. We especially thank Johannes Köster for help creating Snakemake workflows. The project was funded by the Kurt Eberhard Bode Foundation within the Deutsches Stiftungszentrum and by the Science Support Centre of the University of Duisburg-Essen (programme for the promotion of excellent early career researchers).

## Data Accessibility

ddRAD data for *G. fossarum* and *D. gonocephala* will be available at NCBI BioProject upon publication. ddRAD data for *A. fluviatilis* I, II and III will be available at NCBI BioProject with accession number PRJNA389679.

## Author Contributions section

MW, HW and FL designed the study. MW and HW collected the samples and wrote the scripts for ddRAD data analyses. MW performed the laboratory work and data analysis. MW and FL interpreted the data. MW wrote the manuscript and all authors contributed to the final version of the manuscript.

