## Appendix S1 for "Individual small in-stream barriers contribute little to strong local population genetic structure five strictly aquatic macroinvertebrate taxa"

### Appendix S1. Details on lab work and data analyses

#### ddRAD lab workflow

After RNA digestion, the DNA concentration was measured using the Qubit dsDNA BR Assay Kit (Life Technologies, Carlsbad, CA, USA). Depending on the concentration, 120 - 800 ng DNA was used for double digestion. DNA of *D. gonocephala* and *A. fluviatilis* was double-digested with the FastDigest restriction enzymes *Csp6I* and *PstI*, whereas for *G. fossarum SdaI* was used instead of *PstI* to account for the larger genome size (all enzymes ThermoFisher Scientific, Waltham, MA, USA). Expected cut frequencies of the restriction enzymes, needed to calculate the amount of adapters during ligation (Peterson, Weber, Kay, Fisher, & Hoekstra, 2012), were estimated by *in silico* digestion using the script genomecut.pl (Rozenberg, https://github.com/evoeco/radtools/). For *D. gonocephala* and *A. fluviatilis*, the closest available reference genomes were used for estimation (*Schmidtea mediterranea*, GenBank accession number: ASXR00000000.1; *Biomphalaria glabrata* NCBI accession number: APKA00000000.1.). *In silico* digestion resulted in average cut frequencies of 381 bp and 306 bp for *Csp6I,* and 18,416 bp and 9785 bp for *PstI* for *D. gonocephala* and *A. fluviatilis*, respectively. For *G. fossarum,* estimates were based on a standard shotgun genomic library (Macher, Leese, Weigand, & Rozenberg, 2017). Average cut frequencies were 279 bp for *Csp6I* and 178,891 bp for *SdaI*. Adapter ligation was followed by a first size selection using SPRIselect magnetic beads (Beckman Coulter, Brea, CA, USA) to remove excess adapters. As a next step, fragments containing both adapters were amplified by PCR with adapter-specific primers using 14 – 18 cycles. PCR products were purified with AmpureXP magnetic beads (Beckman Coulter), followed by a double size selection with SPRIselect. The DNA concentration after PCR was measured using the Qubit dsDNA HS Assay Kit (Life Technologies; *G. fossarum* library 1) or with the High Sensitivity NGS Fragment Analysis Kit (Advanced Analytical, Ankeny, IA, USA; all other libraries). All samples for each library were pooled in equimolar concentrations, followed by a final precise size selection either using the LabChip XTE (PerkinElmer) with a size range of 308-461 bp (*G. fossarum*: library 1-3, *D. gonocephala*: library 1-3) or E-Gel Power Snap Electrophoresis Device using the E-Gel SizeSelect II gels (2%) with a size range of 300 – 460 bp (*G. fossarum* library 4, *D. gonocephala* library 4, *A. fluviatilis* library 1 and 2). Final libraries were sequenced on an Illumina HiSeq 2500 sequencer using 125 bp paired-end reads (GATC Biotech AG/ Eurofins Genomics Europe Sequencing GmbH; Constance, Germany). Details for ddRAD library preparation for each sample are given in Supporting Information Table S2.

**Stacks parameter validation**

According to the guidelines described by Paris et al. (2017), the parameters M and n were varied between Stacks runs, fixing the parameter m (minimum number of raw reads required to form a stack or putative allele) fixed at 3, as this appeared to be a reasonable setting for many species. M (number of mismatches allowed between stacks for merging them into a putative locus) was varied between 1 and 4 for *A. fluviatilis* and *D. gonocephala*, to account for the polyploid genome of *A. fluviatilis* and possible polyploids in *D. gonocephala*. For *G. fossarum*, this parameter was varied between 2 and 5 to account for high expected levels of polymorphisms. N (number of mismatches allowed to align secondary reads to putative loci) was set to M+2. The number of mismatches allowed between stacks during construction of the catalogue (n) was equal to M or M+1 to account for high the expected diversity within species. For all *Ancylus* species, better results were obtained when Stacks runs included all three species.

**Parameter evaluation**

The aim of the first workflow (step 4, Fig. 3) was to evaluate the eight Stacks parameter and locus filtering settings. The first altered filtering setting was the maximum number of SNPs allowed per locus (1 to 5/7/9 or 12), of which only one was used in further analysis. Additionally, two settings for minor allele frequency (1% or 2%) and the minimum percentage of individuals required for a locus (lim 90% or lim 95%) were tested. To compare results for different Stacks settings, basic population genetic statistics, i.e., observed heterozygosity ($H_O$), within population gene diversity ($H_S$), overall gene diversity ($H_T$), overall $F_{ST}$ and $F_{IS}$ (after Weir & Cockerham (1984)) were calculated using the R-package hierfstat (Goudet, 2005) in R v. 3.3.2 (R Core Team, 2015). Further, principal component analyses (PCAs; Patterson, Price, & Reich, 2006) were performed and individual ancestry coefficients were estimated based on sparse nonnegative matrix factorization algorithms (sNMF; Frichot, Mathieu, Trouillon, Bouchard, & François, 2014) with the R-package LEA (Frichot & François, 2015) to identify the number of genetic clusters per species. Both methods are suitable for polyploid and mixed ploidy data as they do not make population genetic assumptions, like Hardy-Weinberg equilibrium (Dufresne, Stift, Vergilino, & Mable, 2014; Frichot et al., 2014). For the sNMF analyses, k = 1-15 clusters were tested, initially with ten replicates and 30,000 iterations per replicate. For the selection of the best replicate and most probable number of clusters (K), cross-entropy values were compared between replicates and clusters. Based on the sNMF results, individuals of *A. fluviatilis* could be assigned to *A. fluviatilis* I, II, and III (Weiss, Weigand, Weigand, & Leese, 2018) and were from here on analysed separately.

**References Appendix S1**


Dufresne, F., Stift, M., Vergilino, R., & Mable, B. K. (2014). Recent progress and challenges in population genetics of polyploid organisms: An overview of current state-of-the-art molecular and statistical tools. *Molecular Ecology*, *23*(1), 40–69. doi: 10.1111/mec.12581

Frichot, E., & François, O. (2015). LEA: An R package for landscape and ecological association studies. *Methods in Ecology and Evolution*, *6*(8), 925–929. doi: 10.1111/2041-210X.12382

Frichot, E., Mathieu, F., Trouillon, T., Bouchard, G., & François, O. (2014). Fast and Efficient Estimation of Individual Ancestry Coefficients. *Genetics*, *196*(4), 973. doi: 10.1534/genetics.113.160572

Goudet, J. (2005). Hierfstat, a package for r to compute and test hierarchical F-statistics. *Molecular Ecology Notes*, *5*(1), 184–186. doi: 10.1111/j.1471-8286.2004.00828.x

Macher, J. N., Leese, F., Weigand, A. M., & Rozenberg, A. (2017). The complete mitochondrial genome of a cryptic amphipod species from the *Gammarus fossarum* complex. *Mitochondrial DNA Part B*, *2*(1), 17–18. doi: 10.1080/23802359.2016.1275844

Paris, J. R., Stevens, J. R., & Catchen, J. M. (2017). Lost in parameter space: A road map for STACKS. *Methods in Ecology and Evolution*. doi: 10.1111/2041-210X.12775

Patterson, N., Price, A. L., & Reich, D. (2006). Population Structure and Eigenanalysis. *PLoS Genetics*, *2*(12), e190. doi: 10.1371/journal.pgen.0020190

Peterson, B. K., Weber, J. N., Kay, E. H., Fisher, H. S., & Hoekstra, H. E. (2012). Double Digest RADseq: An Inexpensive Method for De Novo SNP Discovery and Genotyping in Model and Non-Model Species. *PLOS ONE*, *7*(5), e37135. doi: 10.1371/journal.pone.0037135

R Core Team. (2015). R: A Language and Environment for Statistical Computing. *R Foundation for Statistical Computing, Vienna, Austria.*, https://www.R-project.org/.

Weir, B., & Cockerham, C. (1984). Estimating F-statistics for the analysis of population structure. *Evolution*, *38*, 1358–1370. doi: 10.2307/2408641

Weiss, M., Weigand, H., Weigand, A. M., & Leese, F. (2018). Genome-wide SNP data reveal cryptic species within cryptic freshwater snail species—The case of the *Ancylus fluviatilis* species complex. *Ecology and Evolution*, *8*(2), 1063–1072. doi: 10.1002/ece3.3706
