## Appendix S2 for "Individual small in-stream barriers contribute little to strong local population genetic structure five strictly aquatic macroinvertebrate taxa"

**Appendix S2.** Nemo .ini files, A) file for random mating species, complete and upstream barrier, B) random mating and additionally reduced downstream migration, C) hermaphrodite species, complete and upstream barriers, D) hermaphrodite species, additionally reduced downstream migration

#### A) NEMO CONFIG FILE FOR RANDOM MATING##
### This is the Nemo config file for random mating simulation, conducted for *Gammarus fossarum* (gf), for complete barrier and upstream barrier simulation

##SIMULATION#
root_dir barrier_gf
logfile logfile_barriergf.log
random_seed 582746
run_mode run
filename %'7[msy10cbmsy10ubmas10cbmas10ubmas15cbmas15ub]'1_fec25_cap%'4'3_ex%2_loc2000

#/
There are three sequential parameters in this init file, "dispersal_matrix", "extinction_proportion", patch_capacity". The filename starts with 'msy10cb', 'msy10ub', 'mas10cb', 'mas10ub', 'mas15cb', or 'mas15ub' for respective dispersal matrix (msy=migration symmetric, mas= migration asymmetric, 10% or 15% are dispersing from source downstream, to next pop rate is the half, asymmetric upstream dispersal half rate (see Fig. 3 for detailed information) patch_capacity (cap), extinction_proportion (ex) are given by their values
/#

replicates 1
generations 250

#### POPULATION ##
patch_number 6 #number of populations
patch_capacity 100 500 1000 2000 #population size

#### LIFE CYCLE EVENTS ##
breed 1
disperse 2
aging 3
extinction 4
save_stats 5
save_files 6

#BREED LCE#
mating_system 1 #random mating
mean_fecundity 25
fecundity_distribution poisson

#DISPERSAL LCE#
### g0 always without barrier (wob), then at g100 different barrier scenarios start (complete (cb) or upstream barrier (ub)) for the different dispersal rates

dispersal_matrix (@g0 {
{0.80625, 0.1, 0.05, 0.025, 0.0125, 0.00625} #msy10wob
{0.1, 0.7125, 0.1, 0.05, 0.025, 0.0125}
{0.05, 0.1, 0.675, 0.1, 0.05, 0.025}
{0.025, 0.05, 0.1, 0.675, 0.1, 0.05}
{0.0125, 0.025, 0.05, 0.1, 0.7125, 0.1}
{0.00625, 0.0125, 0.025, 0.05, 0.1, 0.80625}}, \
@g100 {
{0.85, 0.1, 0.05, 0, 0, 0} #msy10cb
{0.1875, 0.7125, 0.1, 0, 0, 0}
{0.225, 0.1, 0.675, 0, 0, 0}
{0, 0, 0, 0.675, 0.1, 0.225}
{0, 0, 0, 0.1, 0.7125, 0.1875}
{0, 0, 0, 0.05, 0.1, 0.85}}) \
\

```
(@g0 {
{0.80625, 0.1, 0.05, 0.025, 0.0125, 0.00625}                    #msy10wob
{0.1, 0.7125, 0.1, 0.05, 0.025, 0.0125}
{0.05, 0.1, 0.675, 0.1, 0.05, 0.025}
{0.025, 0.05, 0.1, 0.675, 0.1, 0.05}
{0.0125, 0.025, 0.05, 0.1, 0.7125, 0.1}
{0.00625, 0.0125, 0.025, 0.05, 0.1, 0.80625}}, \
@g100 {
{0.80625, 0.1, 0.05, 0.025, 0.0125, 0.00625}                    #msy10ub
{0.1, 0.7125, 0.1, 0.05, 0.025, 0.0125}
{0.05, 0.1, 0.675, 0.1, 0.05, 0.025}
{0, 0, 0, 0.675, 0.1, 0.225}
{0, 0, 0, 0.1, 0.7125, 0.1875}
{0, 0, 0, 0.05, 0.1, 0.85}}) \
\
(@g0 {
{0.80625, 0.1, 0.05, 0.025, 0.0125, 0.00625}                    #mas10wob
{0.05, 0.7625, 0.1, 0.05, 0.025, 0.0125}
{0.025, 0.05, 0.75, 0.1, 0.05, 0.025}
{0.0125, 0.025, 0.05, 0.7625, 0.1, 0.05}
{0.00625, 0.0125, 0.025, 0.05, 0.80625, 0.1}
{0.003125, 0.00625, 0.0125, 0.025, 0.05, 0.903125}}, \
@g100 {
{0.85, 0.1, 0.05, 0, 0, 0}                                      #mas10cb
{0.1375, 0.7625, 0.1, 0, 0, 0}
{0.2, 0.05, 0.75, 0, 0, 0}
{0, 0, 0, 0.7625, 0.1, 0.1375}
{0, 0, 0, 0.05, 0.80625, 0.14375}
{0, 0, 0, 0.025, 0.05, 0.925}}) \
\
(@g0 {
{0.80625, 0.1, 0.05, 0.025, 0.0125, 0.00625}                    #mas10wob
{0.05, 0.7625, 0.1, 0.05, 0.025, 0.0125}
{0.025, 0.05, 0.75, 0.1, 0.05, 0.025}
{0.0125, 0.025, 0.05, 0.7625, 0.1, 0.05}
{0.00625, 0.0125, 0.025, 0.05, 0.80625, 0.1}
{0.003125, 0.00625, 0.0125, 0.025, 0.05, 0.903125}}, \
@g100 {
{0.80625, 0.1, 0.05, 0.025, 0.0125, 0.00625}                    #mas10ub
{0.05, 0.7625, 0.1, 0.05, 0.025, 0.0125}
{0.025, 0.05, 0.75, 0.1, 0.05, 0.025}
{0, 0, 0, 0.7625, 0.1, 0.1375}
{0, 0, 0, 0.05, 0.80625, 0.14375}
{0, 0, 0, 0.025, 0.05, 0.925}}) \
\
(@g0 {
{0.709375, 0.15, 0.075, 0.0375, 0.01875, 0.009375}              #mas15wob
{0.075, 0.64375, 0.15, 0.075, 0.0375, 0.01875}
{0.0375, 0.075, 0.625, 0.15, 0.075, 0.0375}
{0.01875, 0.0375, 0.075, 0.64375, 0.15, 0.075}
{0.009375, 0.01875, 0.0375, 0.075, 0.709375, 0.15}
{0.0046875, 0.009375, 0.01875, 0.0375, 0.075, 0.8546875}}, \
@g100 {
{0.775, 0.15, 0.075, 0, 0, 0}                                   #mas15cb
{0.20625, 0.64375, 0.15, 0, 0, 0}
{0.3, 0.075, 0.625, 0, 0, 0}
{0, 0, 0, 0.64375, 0.15, 0.20625}
{0, 0, 0, 0.075, 0.709375, 0.215625}
{0, 0, 0, 0.0375, 0.075, 0.8875}}) \
\
```

(@g0 {
{0.709375, 0.15, 0.075, 0.0375, 0.01875, 0.009375} #mas15wob
{0.075, 0.64375, 0.15, 0.075, 0.0375, 0.01875}
{0.0375, 0.075, 0.625, 0.15, 0.075, 0.0375}
{0.01875, 0.0375, 0.075, 0.64375, 0.15, 0.075}
{0.009375, 0.01875, 0.0375, 0.075, 0.709375, 0.15}
{0.0046875, 0.009375, 0.01875, 0.0375, 0.075, 0.8546875}}, \
@g100 {
{0.709375, 0.15, 0.075, 0.0375, 0.01875, 0.009375} #mas15ub
{0.075, 0.64375, 0.15, 0.075, 0.0375, 0.01875}
{0.0375, 0.075, 0.625, 0.15, 0.075, 0.0375}
{0, 0, 0, 0.64375, 0.15, 0.20625}
{0, 0, 0, 0.075, 0.709375, 0.215625}
{0, 0, 0, 0.0375, 0.075, 0.8875}})

#EXTINCTION LCE#
extinction_rate 1
extinction_proportion 0.0 0.6 0.8

##OUTPUT##
stat adlt.demography extrate migrants migrants.patch
stat_log_time 10
stat_dir stats_gf

#### NEUTRAL MARKERS ##
ntrl_loci 2000
ntrl_all 2
ntrl_mutation_rate 0.0000001
ntrl_recombination_rate 0.25
ntrl_mutation_model 2

### output marker #
ntrl_save_genotype fstat
ntrl_output_dir genotypes
ntrl_output_logtime 5

#### B) NEMO CONFIG FILE FOR RANDOM MATING##
### This is the Nemo config file for random mating simulation, conducted for *Gammarus fossarum* (gf), for additionally reduced downstream migration

root_dir barrier_gf_reddown
logfile logfile_barriergf_reddown.log
random_seed 582746
run_mode run
filename %'11[mas15ubrd01mas15ubrd02mas15ubrd05]'1_fec25_cap%'4'3_ex%2_loc2000

#/
There are three sequential parameters in this init file, "dispersal_matrix", "extinction_proportion", "patch_capacity". The filename starts with 'mas15ubrd01', 'mas15ubrd02', 'mas15ubrd05' dispersal matrix (mas= migration asymmetric, 15% are dispersing from source downstream to next pop rate is the half, asymmetric upstream half rate. Complete upstream barrier and 1, 2 or 5% dispersal downstream over the barrier (then divided by 2) → reduced downstream (rd), patch_capacity (cap), extinction_proportion (ex) are given by their values
/#

replicates 1
generations 250

#### POPULATION ##
patch_number 6
patch_capacity 500 1000 2000

```
## LIFE CYCLE EVENTS ##
breed          1
disperse       2
aging          3
extinction     4
save_stats     5
save_files     6

#BREED LCE#
mating_system 1                  #random mating
mean_fecundity 25
fecundity_distribution poisson

#DISPERSAL LCE#
# g0 always without barrier, then at g100 different barrier scenarios start (upstream barrier + reduced downstream migration) for the different dispersal rates
dispersal_matrix (@g0 {
{0.709375, 0.15, 0.075, 0.0375, 0.01875, 0.009375}          #mas15wob
{0.075, 0.64375, 0.15, 0.075, 0.0375, 0.01875}
{0.0375, 0.075, 0.625, 0.15, 0.075, 0.0375}
{0.01875, 0.0375, 0.075, 0.64375, 0.15, 0.075}
{0.009375, 0.01875, 0.0375, 0.075, 0.709375, 0.15}
{0.0046875, 0.009375, 0.01875, 0.0375, 0.075, 0.8546875}}, \
@g100 {
{0.7575, 0.15, 0.075, 0.01, 0.005, 0.0025}                  #mas15ubrd01
{0.18875, 0.64375, 0.15, 0.01, 0.005, 0.0025}
{0.2825, 0.075, 0.625, 0.01, 0.005, 0.0025}
{0, 0, 0, 0.64375, 0.15, 0.20625}
{0, 0, 0, 0.075, 0.709375, 0.215625}
{0, 0, 0, 0.0375, 0.075, 0.8875}}) \
\
(@g0 {
{0.709375, 0.15, 0.075, 0.0375, 0.01875, 0.009375}          #mas15wob
{0.075, 0.64375, 0.15, 0.075, 0.0375, 0.01875}
{0.0375, 0.075, 0.625, 0.15, 0.075, 0.0375}
{0.01875, 0.0375, 0.075, 0.64375, 0.15, 0.075}
{0.009375, 0.01875, 0.0375, 0.075, 0.709375, 0.15}
{0.0046875, 0.009375, 0.01875, 0.0375, 0.075, 0.8546875}}, \
@g100 {
{0.73125, 0.15, 0.075, 0.025, 0.0125, 0.00625}              #mas15ubrd02
{0.1625, 0.64375, 0.15, 0.025, 0.0125, 0.00625}
{0.25625, 0.075, 0.625, 0.025, 0.0125, 0.00625}
{0, 0, 0, 0.64375, 0.15, 0.20625}
{0, 0, 0, 0.075, 0.709375, 0.215625}
{0, 0, 0, 0.0375, 0.075, 0.8875}}) \
\
(@g0 {
{0.709375, 0.15, 0.075, 0.0375, 0.01875, 0.009375}          #mas15wob
{0.075, 0.64375, 0.15, 0.075, 0.0375, 0.01875}
{0.0375, 0.075, 0.625, 0.15, 0.075, 0.0375}
{0.01875, 0.0375, 0.075, 0.64375, 0.15, 0.075}
{0.009375, 0.01875, 0.0375, 0.075, 0.709375, 0.15}
{0.0046875, 0.009375, 0.01875, 0.0375, 0.075, 0.8546875}}, \
@g100 {
{0.6875, 0.15, 0.075, 0.05, 0.025, 0.0125}                  #mas15ubrd05
{0.11875, 0.64375, 0.15, 0.05, 0.025, 0.0125}
{0.2125, 0.075, 0.625, 0.05, 0.025, 0.0125}
{0, 0, 0, 0.64375, 0.15, 0.20625}
{0, 0, 0, 0.075, 0.709375, 0.215625}
{0, 0, 0, 0.0375, 0.075, 0.8875}})
```

#EXTINCTION LCE#
extinction_rate 1
extinction_proportion 0.0 0.6

##OUTPUT##
stat adlt.demography extrate migrants migrants.patch
stat_log_time 10
stat_dir stats_gf

#### NEUTRAL MARKERS ##
ntrl_loci 2000
ntrl_all 2
ntrl_mutation_rate 0.0000001
ntrl_recombination_rate 0.25
ntrl_mutation_model 2

### output marker #
ntrl_save_genotype fstat
ntrl_output_dir genotypes
ntrl_output_logtime 5

#### C) NEMO CONFIG FILE FOR HERMAPHRODITES##
### This is the Nemo config file for hermaphroditic species *D. gonocephala* and *A. fluviatilis*, for complete barrier, and upstream barrier simulation

##SIMULATION#
root_dir barrier_herm
logfile logfile_barrier_herm.log
random_seed 582746
run_mode run
filename herm_%'7[msy10cbmsy10ubmas10cbmas10ubmas15cbmas15ub]'1_fec25_cap%'4'4_ex%2_mp%3

#/
There are four sequential parameters in this init file, "dispersal_matrix", "extinction_proportion", "mating_proportion", and "patch_capacity". The filename starts with 'msy10cb', 'msy10ub', 'mas10cb', 'mas10ub', 'mas15cb', or 'mas15ub', dispersal matrix (msy=migration symmetric, mas= migration asymmetric, 10 or 15% are dispersing from source downstream, to next pop rate is the half, asymmetric upstream half rate
→patch_capacity (cap), extinction_proportion (ex) and mating_proportion (mp) are given by their values
/#

replicates 1
generations 250

#### POPULATION ##
patch_number 6
patch_capacity 50 100 200 500 1000 2000

#### LIFE CYCLE EVENTS ##
breed 1
disperse 2
aging 3
extinction 4
save_stats 5
save_files 6

#BREED LCE#
mating_system 4 #hermaphrodite
mating_proportion 0.0 0.3 0.6 0.9 1.0
mean_fecundity 25
fecundity_distribution poisson

```
#DISPERSAL LCE#
# g0 always without barrier, then at g100different barrier scenarios start (complete or upstream barrier) for the different dispersal rates
dispersal_matrix (@g0 {
{0.80625, 0.1, 0.05, 0.025, 0.0125, 0.00625}                    #msy10wob
{0.1, 0.7125, 0.1, 0.05, 0.025, 0.0125}
{0.05, 0.1, 0.675, 0.1, 0.05, 0.025}
{0.025, 0.05, 0.1, 0.675, 0.1, 0.05}
{0.0125, 0.025, 0.05, 0.1, 0.7125, 0.1}
{0.00625, 0.0125, 0.025, 0.05, 0.1, 0.80625}}, \
@g100 {
{0.85, 0.1, 0.05, 0, 0, 0}                                      #msy10cb
{0.1875, 0.7125, 0.1, 0, 0, 0}
{0.225, 0.1, 0.675, 0, 0, 0}
{0, 0, 0, 0.675, 0.1, 0.225}
{0, 0, 0, 0.1, 0.7125, 0.1875}
{0, 0, 0, 0.05, 0.1, 0.85}}) \
\
(@g0 {
{0.80625, 0.1, 0.05, 0.025, 0.0125, 0.00625}                    #msy10wob
{0.1, 0.7125, 0.1, 0.05, 0.025, 0.0125}
{0.05, 0.1, 0.675, 0.1, 0.05, 0.025}
{0.025, 0.05, 0.1, 0.675, 0.1, 0.05}
{0.0125, 0.025, 0.05, 0.1, 0.7125, 0.1}
{0.00625, 0.0125, 0.025, 0.05, 0.1, 0.80625}}, \
@g100 {
{0.80625, 0.1, 0.05, 0.025, 0.0125, 0.00625}                    #msy10ub
{0.1, 0.7125, 0.1, 0.05, 0.025, 0.0125}
{0.05, 0.1, 0.675, 0.1, 0.05, 0.025}
{0, 0, 0, 0.675, 0.1, 0.225}
{0, 0, 0, 0.1, 0.7125, 0.1875}
{0, 0, 0, 0.05, 0.1, 0.85}}) \
\
(@g0 {
{0.80625, 0.1, 0.05, 0.025, 0.0125, 0.00625}                    #mas10wob
{0.05, 0.7625, 0.1, 0.05, 0.025, 0.0125}
{0.025, 0.05, 0.75, 0.1, 0.05, 0.025}
{0.0125, 0.025, 0.05, 0.7625, 0.1, 0.05}
{0.00625, 0.0125, 0.025, 0.05, 0.80625, 0.1}
{0.003125, 0.00625, 0.0125, 0.025, 0.05, 0.903125}}, \
@g100 {
{0.85, 0.1, 0.05, 0, 0, 0}                                      #mas10cb
{0.1375, 0.7625, 0.1, 0, 0, 0}
{0.2, 0.05, 0.75, 0, 0, 0}
{0, 0, 0, 0.7625, 0.1, 0.1375}
{0, 0, 0, 0.05, 0.80625, 0.14375}
{0, 0, 0, 0.025, 0.05, 0.925}}) \
\
(@g0 {
{0.80625, 0.1, 0.05, 0.025, 0.0125, 0.00625}                    #mas10wob
{0.05, 0.7625, 0.1, 0.05, 0.025, 0.0125}
{0.025, 0.05, 0.75, 0.1, 0.05, 0.025}
{0.0125, 0.025, 0.05, 0.7625, 0.1, 0.05}
{0.00625, 0.0125, 0.025, 0.05, 0.80625, 0.1}
{0.003125, 0.00625, 0.0125, 0.025, 0.05, 0.903125}}, \
@g100 {
{0.80625, 0.1, 0.05, 0.025, 0.0125, 0.00625}                    #mas10ub
{0.05, 0.7625, 0.1, 0.05, 0.025, 0.0125}
{0.025, 0.05, 0.75, 0.1, 0.05, 0.025}
{0, 0, 0, 0.7625, 0.1, 0.1375}
{0, 0, 0, 0.05, 0.80625, 0.14375}
{0, 0, 0, 0.025, 0.05, 0.925}}) \
```

```
\
(@g0 {
{0.709375, 0.15, 0.075, 0.0375, 0.01875, 0.009375}              #mas15wob
{0.075, 0.64375, 0.15, 0.075, 0.0375, 0.01875}
{0.0375, 0.075, 0.625, 0.15, 0.075, 0.0375}
{0.01875, 0.0375, 0.075, 0.64375, 0.15, 0.075}
{0.009375, 0.01875, 0.0375, 0.075, 0.709375, 0.15}
{0.0046875, 0.009375, 0.01875, 0.0375, 0.075, 0.8546875}}, \
@g100 {
{0.775, 0.15, 0.075, 0, 0, 0}                                   #mas15cb
{0.20625, 0.64375, 0.15, 0, 0, 0}
{0.3, 0.075, 0.625, 0, 0, 0}
{0, 0, 0, 0.64375, 0.15, 0.20625}
{0, 0, 0, 0.075, 0.709375, 0.215625}
{0, 0, 0, 0.0375, 0.075, 0.8875}}) \
\
(@g0 {
{0.709375, 0.15, 0.075, 0.0375, 0.01875, 0.009375}              #mas15wob
{0.075, 0.64375, 0.15, 0.075, 0.0375, 0.01875}
{0.0375, 0.075, 0.625, 0.15, 0.075, 0.0375}
{0.01875, 0.0375, 0.075, 0.64375, 0.15, 0.075}
{0.009375, 0.01875, 0.0375, 0.075, 0.709375, 0.15}
{0.0046875, 0.009375, 0.01875, 0.0375, 0.075, 0.8546875}}, \
@g100 {
{0.709375, 0.15, 0.075, 0.0375, 0.01875, 0.009375}              #mas15ub
{0.075, 0.64375, 0.15, 0.075, 0.0375, 0.01875}
{0.0375, 0.075, 0.625, 0.15, 0.075, 0.0375}
{0, 0, 0, 0.64375, 0.15, 0.20625}
{0, 0, 0, 0.075, 0.709375, 0.215625}
{0, 0, 0, 0.0375, 0.075, 0.8875}})

#EXTINCTION LCE#
extinction_rate 1
extinction_proportion 0.0 0.6 0.8

##OUTPUT##
stat adlt.demography extrate migrants migrants.patch
stat_log_time 10
stat_dir stats_gf

## NEUTRAL MARKERS ##
ntrl_loci 2000
ntrl_all 2
ntrl_mutation_rate 0.0000001
ntrl_recombination_rate 0.25
ntrl_mutation_model 2

# output marker #
ntrl_save_genotype fstat
ntrl_output_dir genotypes
ntrl_output_logtime 5
```

##D) NEMO CONFIG FILE FOR HERMAPHRODITES##
### This is the Nemo config file for hermaphroditic species *D. gonocephala* and *A. fluviatilis*, for additionally reduced downstream migration

##SIMULATION#
root_dir barrier_herm_reddown
logfile logfile_barrier_herm_reddown.log
random_seed 582746
run_mode overwrite
filename %'11[mas15ubrd01mas15ubrd02mas15ubrd05]'1_fec25_cap%'4'4_ex%2_loc2000_mp%3

#/
There are four sequential parameters in this init file, "dispersal_matrix", "extinction_proportion", "mating_proportion", and "patch_capacity". The filename starts with 'mas15ubrd01', 'mas15ubrd02', 'mas15ubrd05' dispersal matrix (mas= migration asymmetric, 15% are dispersing from source downstream to next pop rate is the half, asymmetric upstream half rate. Complete upstream barrier and 1, 2 or 5% dispersal downstream over the barrier (then divided by 2) →reduced downstream (rd), patch_capacity (cap), extinction_proportion (ex) and mating_proportion (mp) are given by their values
/#

replicates 1
generations 250

#### POPULATION ##
patch_number 6
patch_capacity 500 1000 2000

#### LIFE CYCLE EVENTS ##
breed 1
disperse 2
aging 3
extinction 4
save_stats 5
save_files 6

#BREED LCE#
mating_system 4 #hermaphrodite
mating_proportion 0.3 0.6
mean_fecundity 25
fecundity_distribution poisson

#DISPERSAL LCE#
### g0 always without barrier, then at g100 different barrier scenarios start (upstream barrier + reduced downstream migration) for the different dispersal rates
dispersal_matrix (@g0 {
{0.709375, 0.15, 0.075, 0.0375, 0.01875, 0.009375} #mas15wob
{0.075, 0.64375, 0.15, 0.075, 0.0375, 0.01875}
{0.0375, 0.075, 0.625, 0.15, 0.075, 0.0375}
{0.01875, 0.0375, 0.075, 0.64375, 0.15, 0.075}
{0.009375, 0.01875, 0.0375, 0.075, 0.709375, 0.15}
{0.0046875, 0.009375, 0.01875, 0.0375, 0.075, 0.8546875}}, \
@g100 {
{0.7575, 0.15, 0.075, 0.01, 0.005, 0.0025} #mas15ubrd01
{0.18875, 0.64375, 0.15, 0.01, 0.005, 0.0025}
{0.2825, 0.075, 0.625, 0.01, 0.005, 0.0025}
{0, 0, 0, 0.64375, 0.15, 0.20625}
{0, 0, 0, 0.075, 0.709375, 0.215625}
{0, 0, 0, 0.0375, 0.075, 0.8875}}) \
\

```
(@g0 {
{0.709375, 0.15, 0.075, 0.0375, 0.01875, 0.009375}              #mas15wob
{0.075, 0.64375, 0.15, 0.075, 0.0375, 0.01875}
{0.0375, 0.075, 0.625, 0.15, 0.075, 0.0375}
{0.01875, 0.0375, 0.075, 0.64375, 0.15, 0.075}
{0.009375, 0.01875, 0.0375, 0.075, 0.709375, 0.15}
{0.0046875, 0.009375, 0.01875, 0.0375, 0.075, 0.8546875}}, \
@g100 {
{0.73125, 0.15, 0.075, 0.025, 0.0125, 0.00625}                  #mas15ubrd02
{0.1625, 0.64375, 0.15, 0.025, 0.0125, 0.00625}
{0.25625, 0.075, 0.625, 0.025, 0.0125, 0.00625}
{0, 0, 0, 0.64375, 0.15, 0.20625}
{0, 0, 0, 0.075, 0.709375, 0.215625}
{0, 0, 0, 0.0375, 0.075, 0.8875}}) \
\
(@g0 {
{0.709375, 0.15, 0.075, 0.0375, 0.01875, 0.009375}              #mas15wob
{0.075, 0.64375, 0.15, 0.075, 0.0375, 0.01875}
{0.0375, 0.075, 0.625, 0.15, 0.075, 0.0375}
{0.01875, 0.0375, 0.075, 0.64375, 0.15, 0.075}
{0.009375, 0.01875, 0.0375, 0.075, 0.709375, 0.15}
{0.0046875, 0.009375, 0.01875, 0.0375, 0.075, 0.8546875}}, \
@g100 {
{0.6875, 0.15, 0.075, 0.05, 0.025, 0.0125}                      #mas15ubrd05
{0.11875, 0.64375, 0.15, 0.05, 0.025, 0.0125}
{0.2125, 0.075, 0.625, 0.05, 0.025, 0.0125}
{0, 0, 0, 0.64375, 0.15, 0.20625}
{0, 0, 0, 0.075, 0.709375, 0.215625}
{0, 0, 0, 0.0375, 0.075, 0.8875}})

#EXTINCTION LCE#
extinction_rate 1
extinction_proportion 0.0 0.6

##OUTPUT##
stat adlt.demography extrate migrants migrants.patch
stat_log_time 10
stat_dir stats_gf

## NEUTRAL MARKERS ##
ntrl_loci 2000
ntrl_all 2
ntrl_mutation_rate 0.0000001
ntrl_recombination_rate 0.25
ntrl_mutation_model 2

# output marker #
# ntrl_save_freq locus
ntrl_save_genotype fstat
ntrl_output_dir genotypes
ntrl_output_logtime 5
```
