## Supplementary figures and images for "Individual small in-stream barriers contribute little to strong local population genetic structure five strictly aquatic macroinvertebrate taxa"

### Figure S1

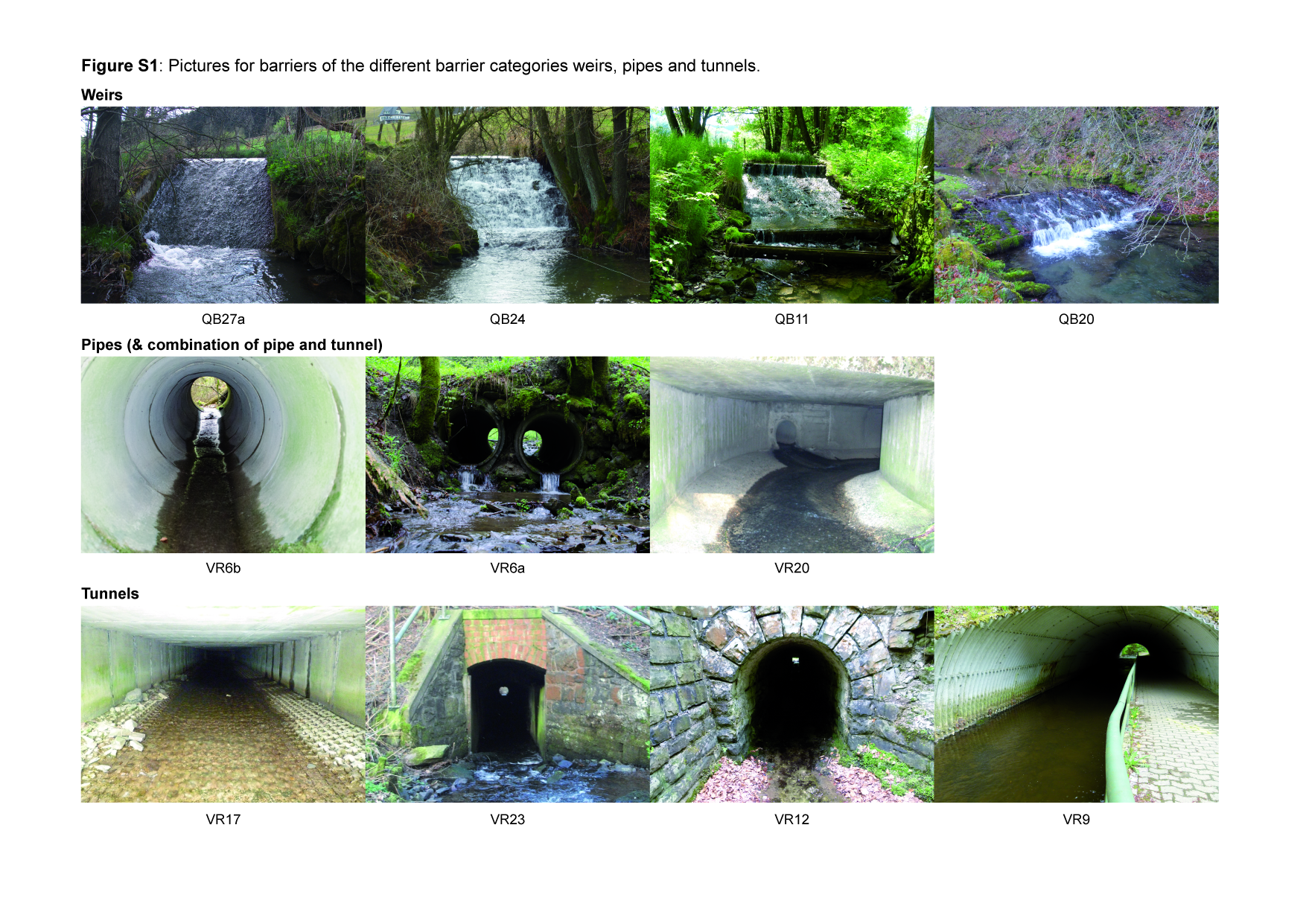
