## Supplementary material for "Individual small in-stream barriers contribute little to strong local population genetic structure five strictly aquatic macroinvertebrate taxa": Figure S2

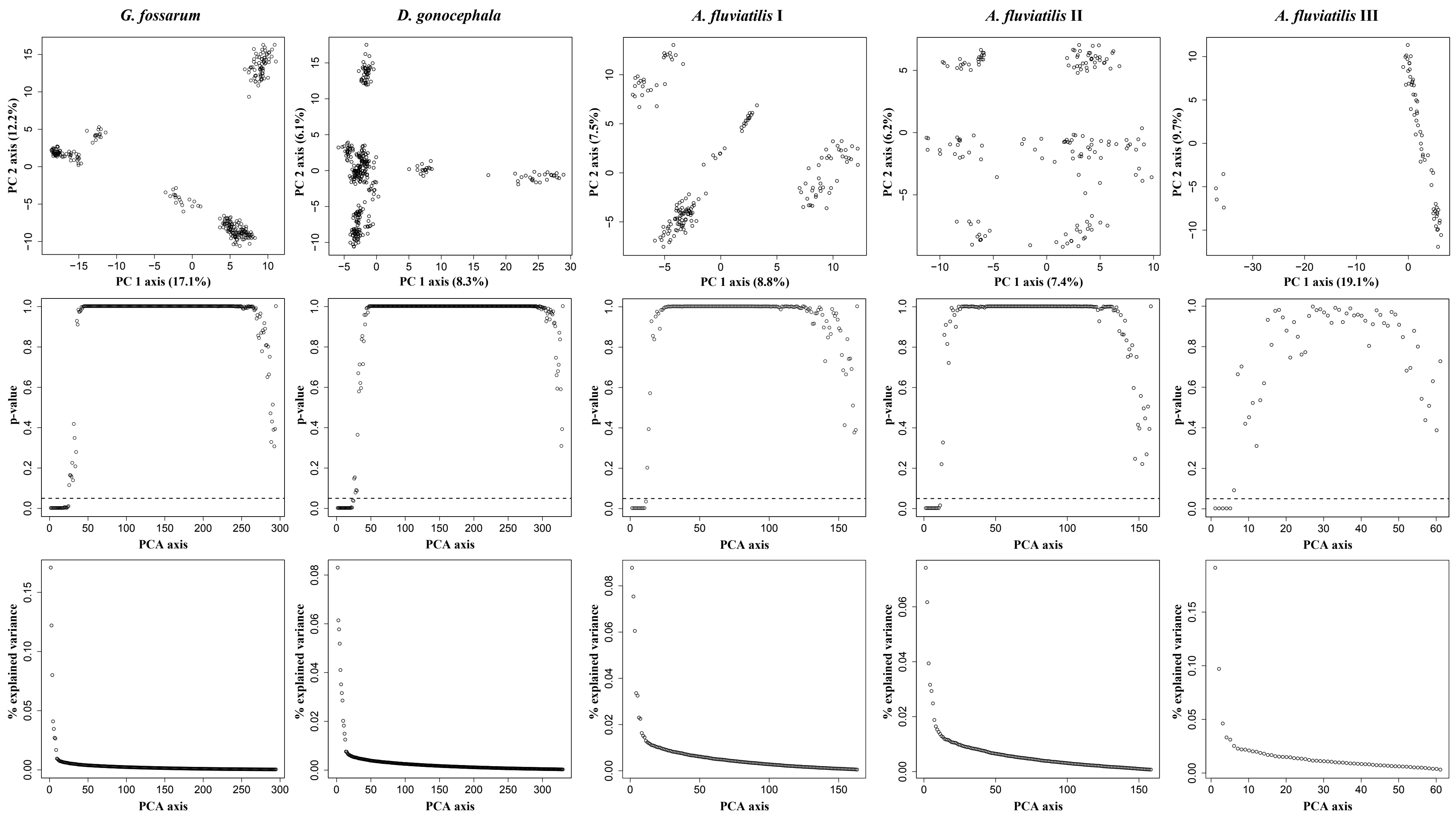


**Figure S2.** Principle component analysis for all species (columns). Scatterplots in the first row show the first two principal components (PCs), percentage of explained variance by the respective axes is given in brackets. In the second row, p-values for all axes for the different species are shown with points below the vertical dashed line indicating significance of axes ($p < 0.05$). Plots in the third row show the percentage of variance explained by each axis.
