## Supplementary material for "Individual small in-stream barriers contribute little to strong local population genetic structure five strictly aquatic macroinvertebrate taxa": Figure S3

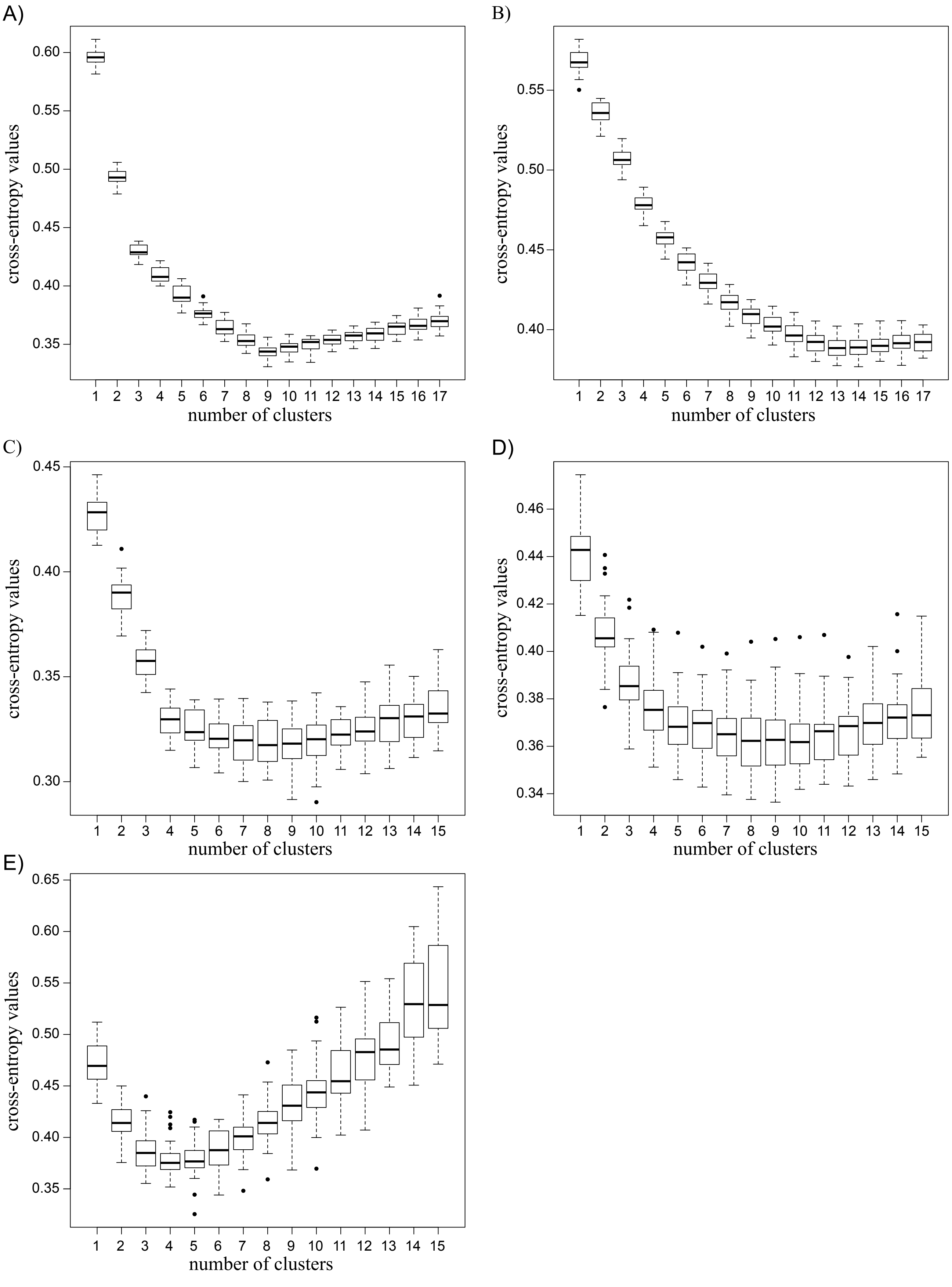


**Figure S3.** Standard boxplots of cross-entropy values (30 repeats) of sNMF analysis for final ddRAD datasets for the different taxa. A) *G. fossarum*, B) *D. gonocephala,* C-E) *A. fluviatilis* I, II and III.
