## Supplementary material for "Individual small in-stream barriers contribute little to strong local population genetic structure five strictly aquatic macroinvertebrate taxa": Figure S4

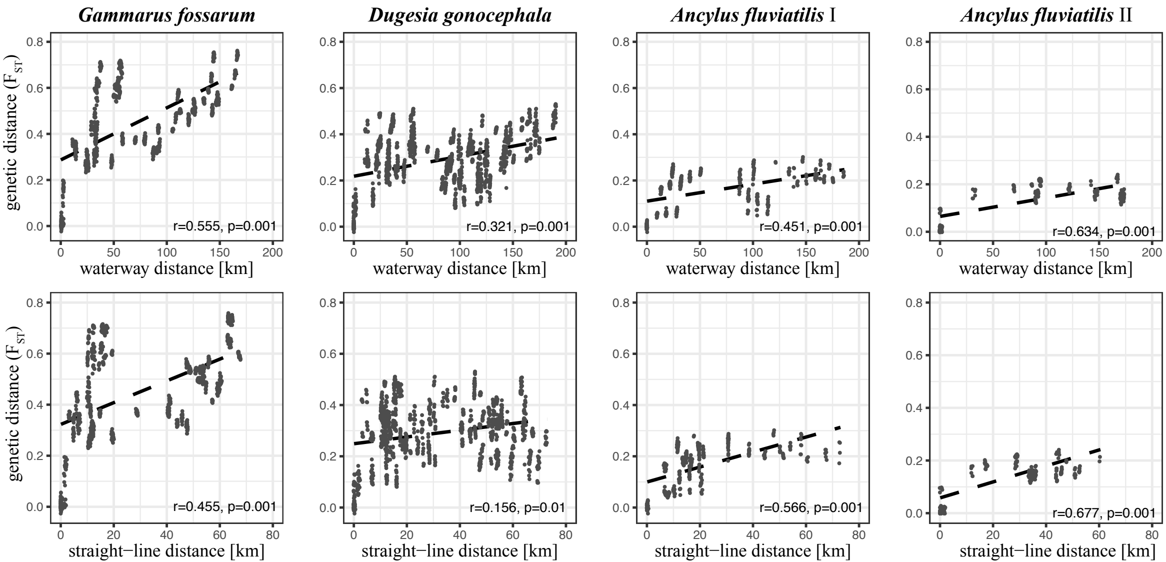


**Figure S4.** Correlation between pairwise genetic distance ($F_{ST}$) among single sampling sites and geographic distances (first row waterway distance; second row straight-line distance) for *G. fossarum*, *D. gonocephala*, *A. fluviatilis* I and *A. fluviatilis* II (different columns).
