## Supplementary material for "Individual small in-stream barriers contribute little to strong local population genetic structure five strictly aquatic macroinvertebrate taxa": Figure S5

| *G. fossarum* (*Gf*) | *D. gonocephala* (*Dg*) | *A. fluviatilis* (*Af*) I, II, III | Summary |
|---|---|---|---|
| | QB11) 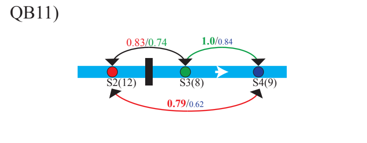 | *Af* II QB11) 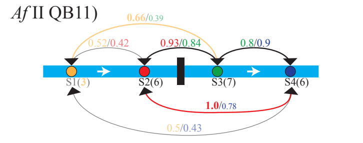 | ***Dg***: migration stronger in downstream (*ds*) than in upstream (*us*) direction; *ds* & *us* migration lower across barrier than without barrier<br>***Af* II**: tendency for stronger *ds* migration, migration from/to S1 reduced, but only n = 3; excluding S1: *ds* migration stronger across barrier, *us* lower across barrier |
| QB12) 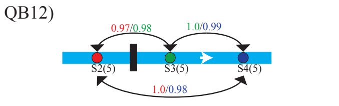 | | *Af* I QB12) 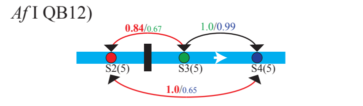 | ***Gf***: generally strong migration, no barrier effect detectable<br>***Af* I**: tendency for stronger *ds* migration; *us* lower across barrier |
| QB17) 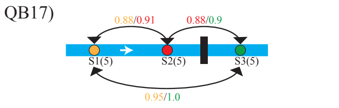 | QB17) 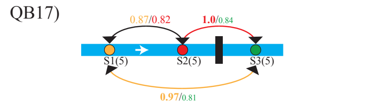 | *Af* II QB17) 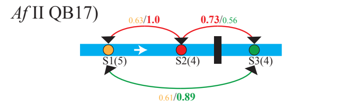 | ***Gf***: generally strong migration, no barrier effect detectable<br>***Dg***: migration stronger in *ds* direction; *ds* migration stronger across barrier, *us* no pattern<br>***Af* II**: *ds* from S1 reduced; *us* lower across barrier |
| | QB20) 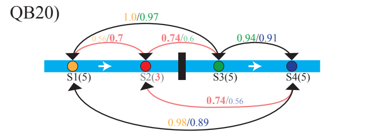 | *Af* I QB20) 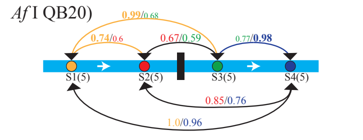 | ***Dg***: migration from/to S2 reduced (n = 3); no barrier effect detectable<br>***Af* I**: tendency for stronger *ds* migration; lowest migration in both directions directly across barrier, but no overall barrier effect detectable |
| QB22) 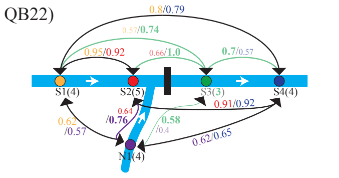 | QB22) 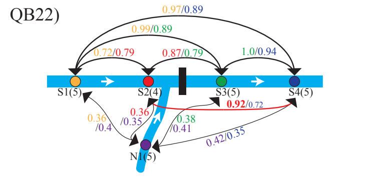 | *Af* I QB22) 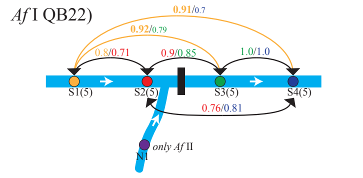 | ***Gf***: migration from/to N1 reduced, no barrier effect detectable<br>***Dg***: migration from/to N1 strongly reduced; no barrier effect detectable<br>***Af* I**: at N1 only *Af* II found, tendency for higher *ds* migration, no barrier effect detectable |
| QB23) 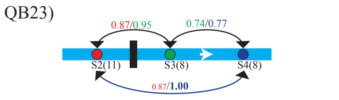 | QB23) 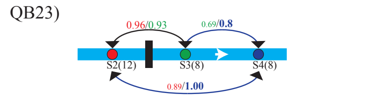 | *Af* II QB23) 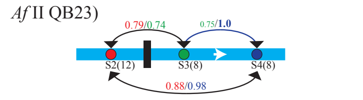 | ***Gf***: tendency for stronger *us* migration; *ds*/*us* migration lowest among reference sites; ***Dg*** similar to *Gf*<br>***Af* II**: tendency for stronger *us* migration, no barrier effect detectable |
| QB24) 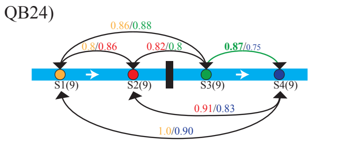 | QB24) 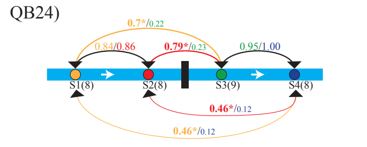 | *Af* I QB24) 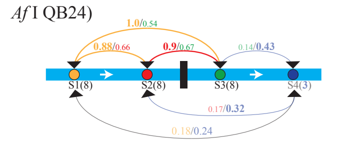 | ***Gf***: no pattern detactable<br>***Dg***: tendency for stronger *ds* migration, *us* migration across barrier significantly reduced, *ds* & *us* migration lower across barrier than among reference sites<br>***Af* I**: migration from/to S4 reduced (n= 3), otherwise stronger *ds* directed migration; no barrier effect detectable |
| QB27) 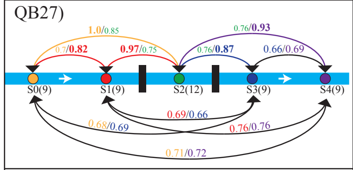 | QB27) 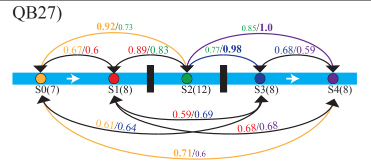 | *Af* I QB27) 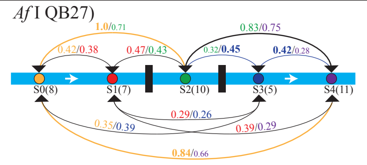 | ***Gf***: no pattern detactable<br>***Dg***: no pattern detactable<br>***Af* I**: generally relatively low migration rates |
| VR6) 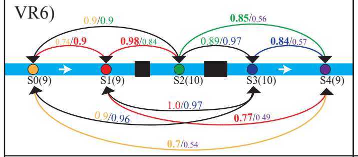 | VR6) 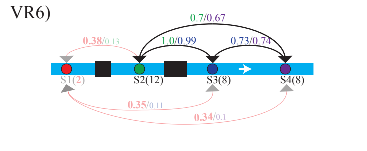 | *Af* III VR6) 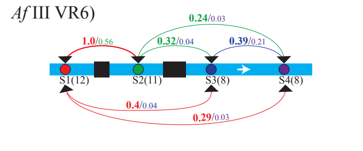 | ***Gf***: migration from S4 reduced, no barrier effect detectable<br>***Dg***: migration from/to S1 strongly reduced, but only n = 2<br>***Af* III**: generelly reduced migration rates, stronger *ds* migration; *us* migration across 2. pipe strongly reduced, 1. pipe stronger *us* migration than among reference sites |
| | VR9) 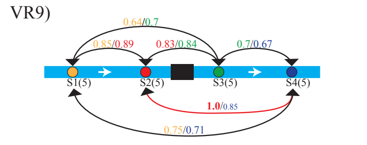 | *Af* I VR9) 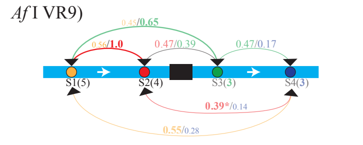 | ***Dg***: no pattern detectable<br>***Af* I**: low sample size below barrier, no clear pattern detectable |
| VR11) 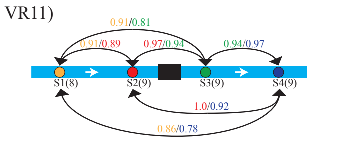 | VR11) 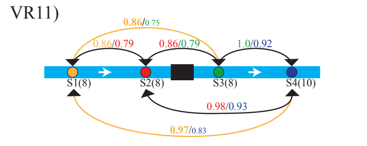 | *Af* II VR11) 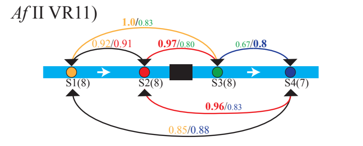 | ***Gf***: no pattern detectable<br>***Dg***: tendency for stronger *ds* migration; no barrier effect detectable<br>***Af* II**: no pattern detectable |
| VR12) 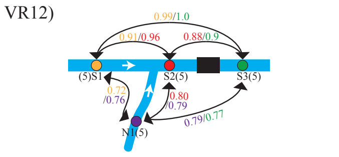 | VR12) 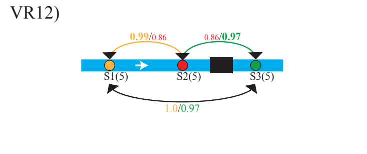 | | ***Gf***: migration rates from/to N1 lower, no barrier effect detectable<br>***Dg***: no pattern detectable |
| VR17) 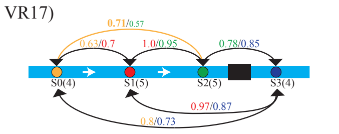 | VR17) 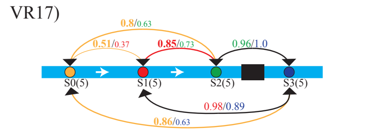 | *Af* III VR17) 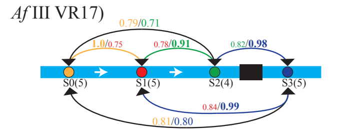 | ***Gf***: no pattern detectable<br>***Dg***: no pattern detectable<br>***Af* III**: no pattern detectable |
| | | *Af* II VR20) 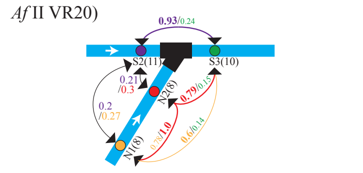 | ***Af* II**: *ds* migration through culvert possible, *us* migration rates through culvert reduced, especially to N sites; strongest reduction of migration rates among sites in streams *us* of the barrier (S2 to N1/N2) |
| VR23) 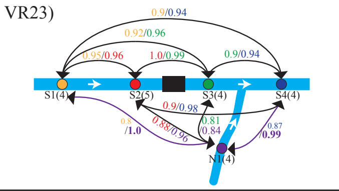 | VR23) 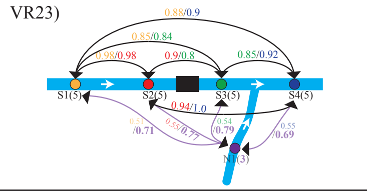 | | ***Gf***: generally high migration rates; no barrier effect detectable<br>***Dg***: migration rates from/to N1 reduced, but n = 3; no barrier effect detectable |

**Figure S5.** Vizualisation of asymmetric migration rates among all single sampling site calculated seperately for each barrier. Rates are colored according to source site and arrows are colored if rates differed more than 0.15 and then according to the direction of the higher rate. Significant asymmetry is indicated by an asterisk. If sample size was ≤ 3, rates are shown in lighter color.
