## Supplementary material for "Individual small in-stream barriers contribute little to strong local population genetic structure five strictly aquatic macroinvertebrate taxa": Figure S6

**Figure S6**: Plots showing changes in population differentiation ($F_{ST}$) among populations over generations for different Nemo simulation scenarios. The barrier, introduced after 100 generations is indicated by a red line. A representative subset of simulations is shown. Results were similar for all migration matrices and are only shown for *asymmetric15* (mas15), but all barrier models (cb = complete barrier, ub = upstream barrier, ubrd01/ 02/ 05 = upstream barrier + reduced downstream migration to 1%/ 2% / or 5%. Simulations without extiction (ex0.0) and an extinction rate of 60% per generation (ex0.6) are shown either for random mating populations or for hermaphrodites (indicated by mp0.3 for mating proportion/selfing rate). In plots on first 3 pages, populations on each site of the barrier were pooled and $F_{ST}$ was calculated among these two populations. In these plots, all simulated patch capacities are shown. In plots on pages 4 - 6, $F_{ST}$ values were calculated among all four single populations and colored according to the presence of a barrier. Here, all barrier models, ex0.0 and ex0.6, random mating populations and hermaphrodites (mp0.3) are shown for patch capacites 500 and 2000.

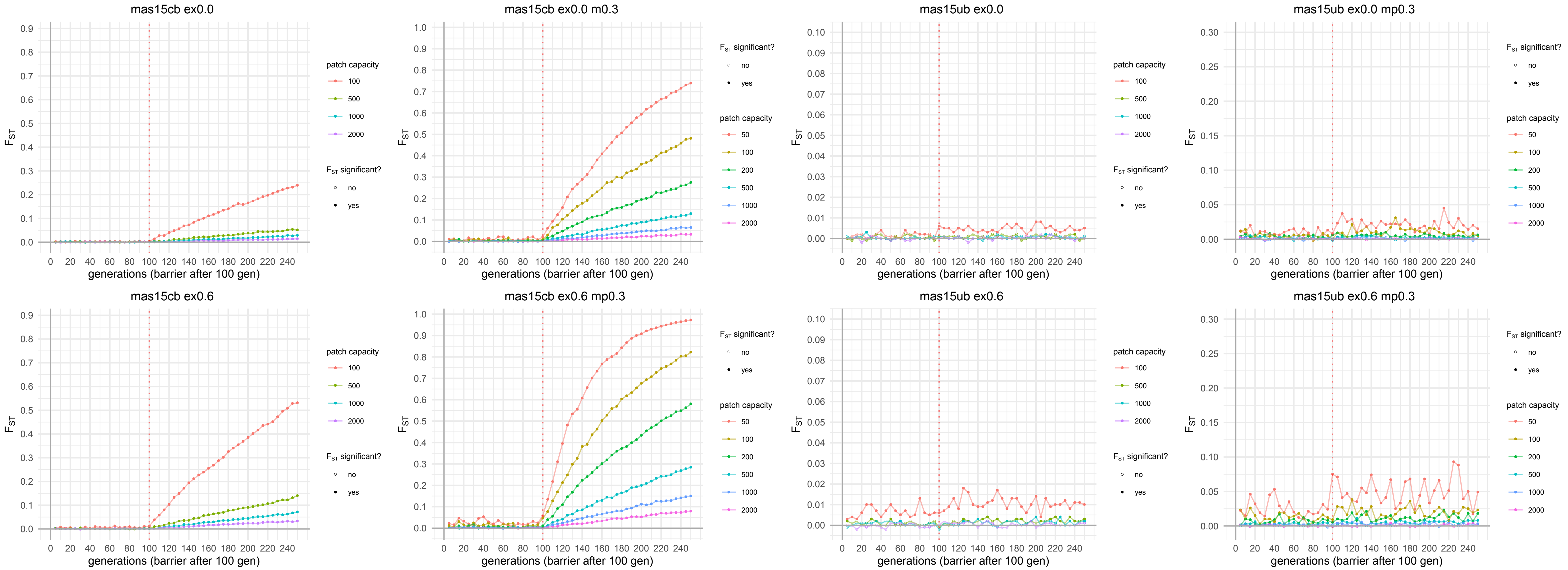

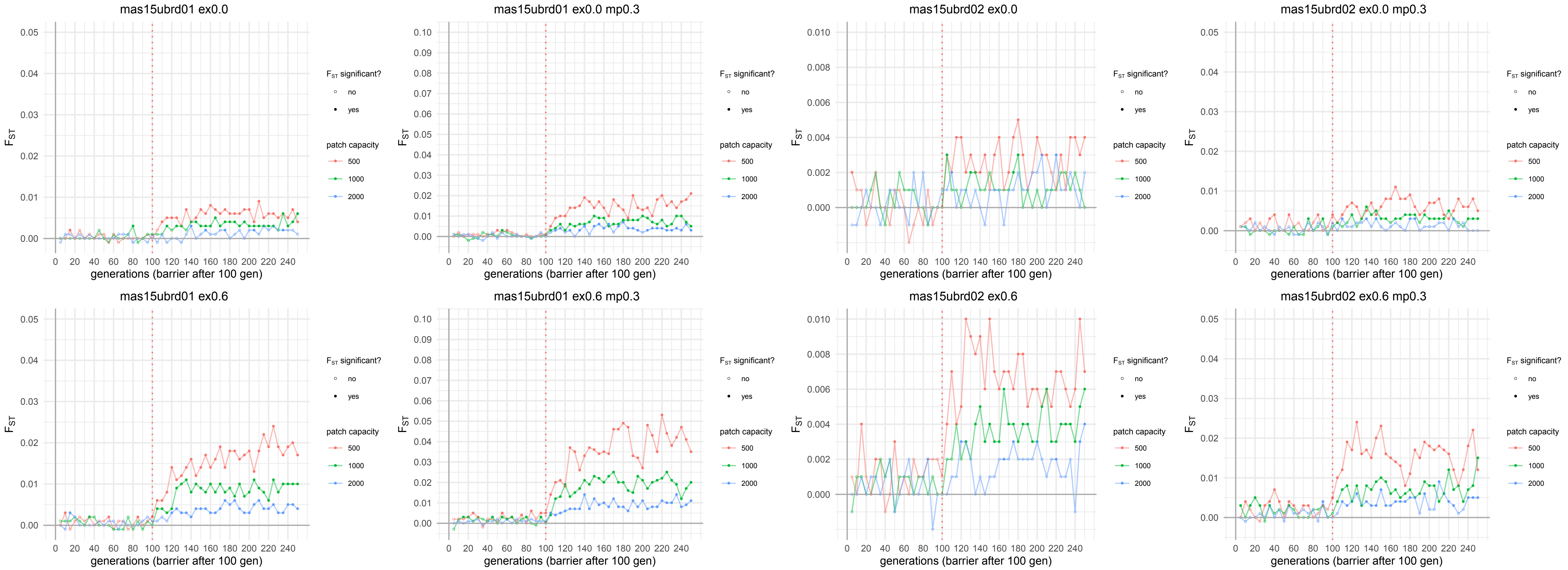
mas15ubrd01 ex0.0
mas15ubrd01 ex0.0 mp0.3
mas15ubrd02 ex0.0
mas15ubrd02 ex0.0 mp0.3
mas15ubrd01 ex0.6
mas15ubrd01 ex0.6 mp0.3
mas15ubrd02 ex0.6
mas15ubrd02 ex0.6 mp0.3
$F_{ST}$
generations (barrier after 100 gen)
$F_{ST}$ significant?
no
yes
patch capacity
500
1000
2000

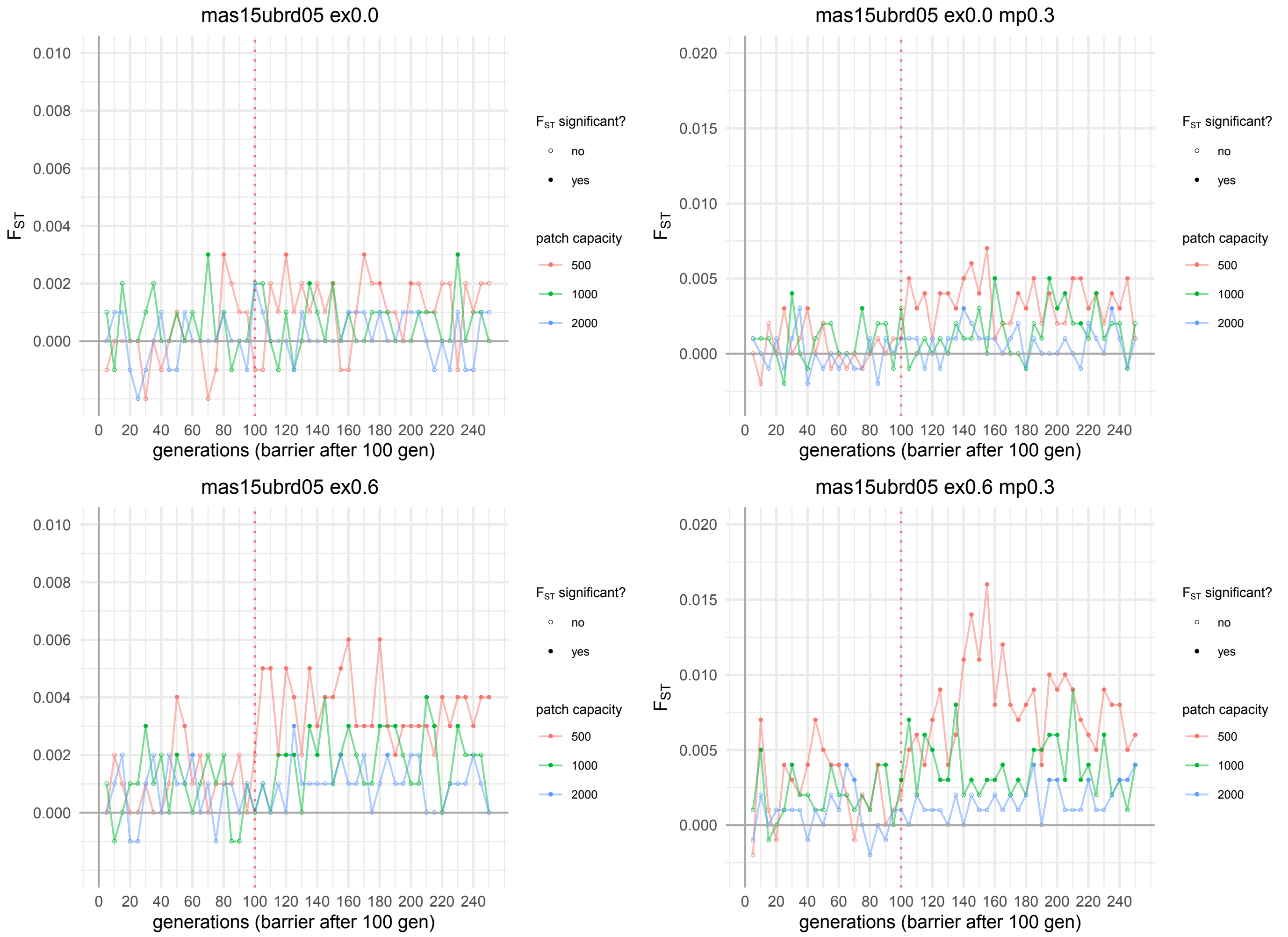

mas15ubrd05 ex0.0
mas15ubrd05 ex0.0 mp0.3
mas15ubrd05 ex0.6
mas15ubrd05 ex0.6 mp0.3
$F_{ST}$
generations (barrier after 100 gen)
0 20 40 60 80 100 120 140 160 180 200 220 240
0.000 0.002 0.004 0.006 0.008 0.010
0.000 0.005 0.010 0.015 0.020
$F_{ST}$ significant?
no
yes
patch capacity
500
1000
2000

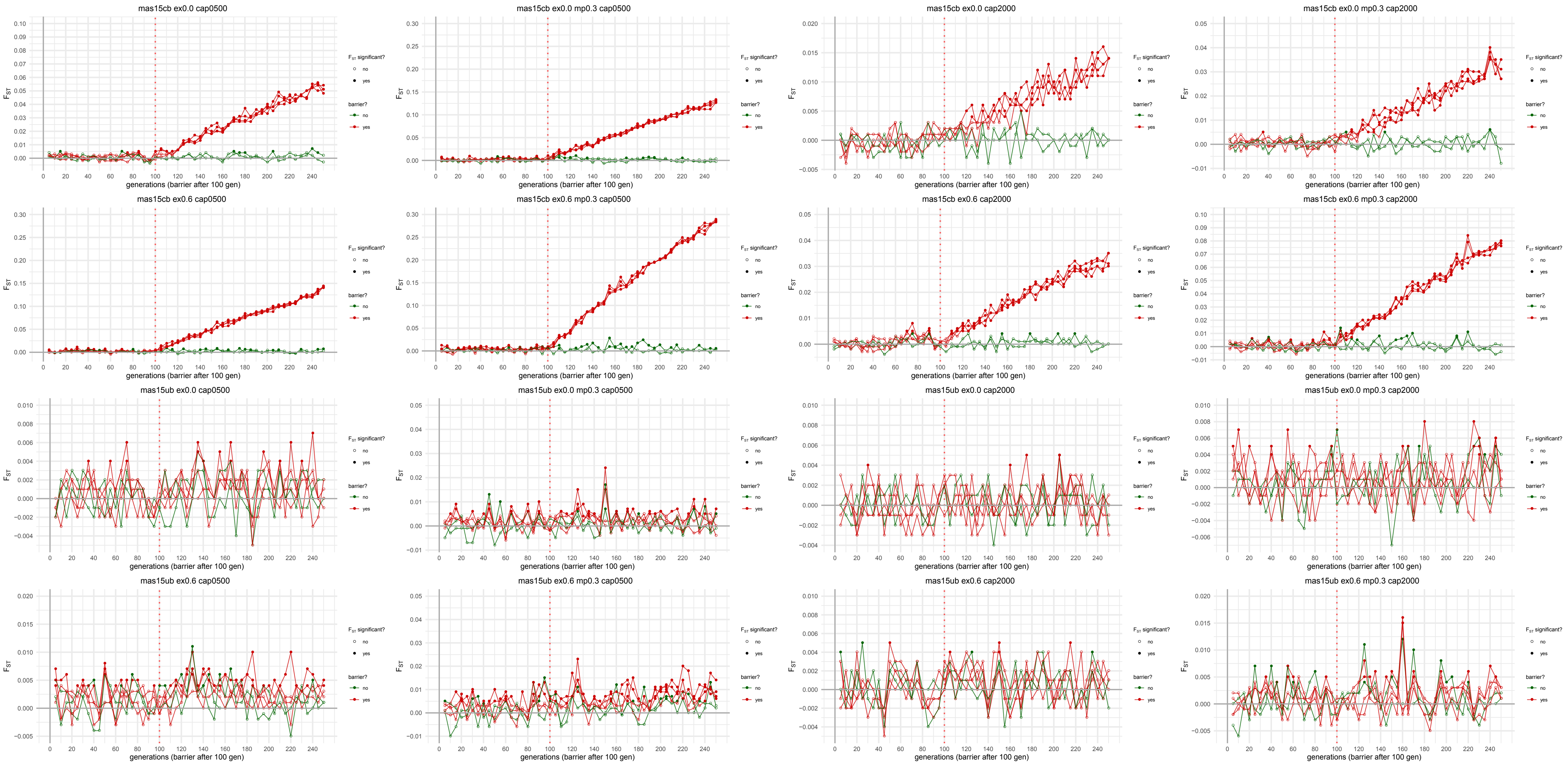
mas15cb ex0.0 cap0500
mas15cb ex0.0 mp0.3 cap0500
mas15cb ex0.0 cap2000
mas15cb ex0.0 mp0.3 cap2000
mas15cb ex0.6 cap0500
mas15cb ex0.6 mp0.3 cap0500
mas15cb ex0.6 cap2000
mas15cb ex0.6 mp0.3 cap2000
mas15ub ex0.0 cap0500
mas15ub ex0.0 mp0.3 cap0500
mas15ub ex0.0 cap2000
mas15ub ex0.0 mp0.3 cap2000
mas15ub ex0.6 cap0500
mas15ub ex0.6 mp0.3 cap0500
mas15ub ex0.6 cap2000
mas15ub ex0.6 mp0.3 cap2000
$F_{ST}$
generations (barrier after 100 gen)
$F_{ST}$ significant?
no
yes
barrier?
no
yes

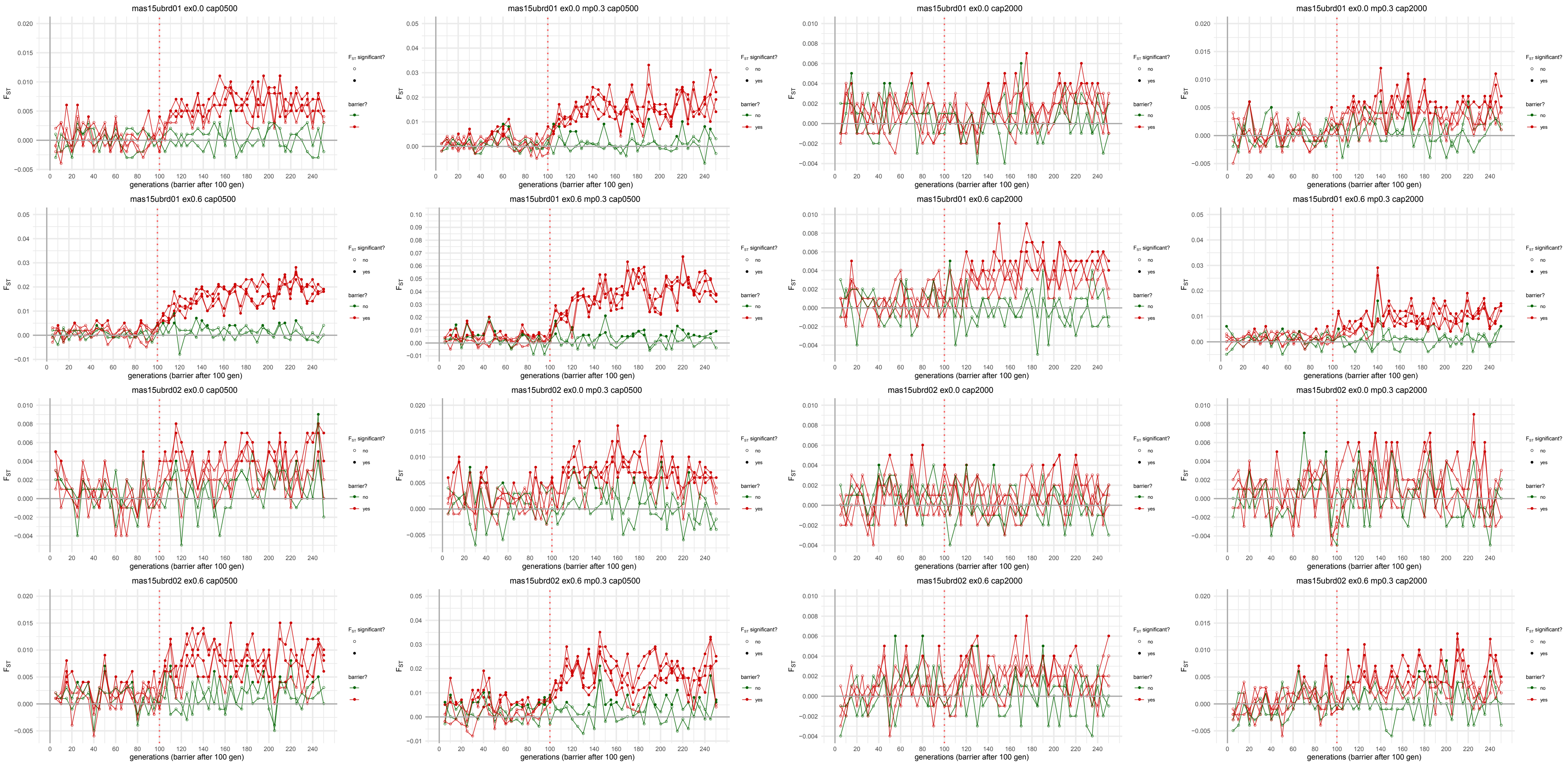
mas15ubrd01 ex0.0 cap0500
mas15ubrd01 ex0.0 mp0.3 cap0500
mas15ubrd01 ex0.0 cap2000
mas15ubrd01 ex0.0 mp0.3 cap2000
mas15ubrd01 ex0.6 cap0500
mas15ubrd01 ex0.6 mp0.3 cap0500
mas15ubrd01 ex0.6 cap2000
mas15ubrd01 ex0.6 mp0.3 cap2000
mas15ubrd02 ex0.0 cap0500
mas15ubrd02 ex0.0 mp0.3 cap0500
mas15ubrd02 ex0.0 cap2000
mas15ubrd02 ex0.0 mp0.3 cap2000
mas15ubrd02 ex0.6 cap0500
mas15ubrd02 ex0.6 mp0.3 cap0500
mas15ubrd02 ex0.6 cap2000
mas15ubrd02 ex0.6 mp0.3 cap2000
$F_{ST}$
generations (barrier after 100 gen)
$F_{ST}$ significant?
no
yes
barrier?
no
yes

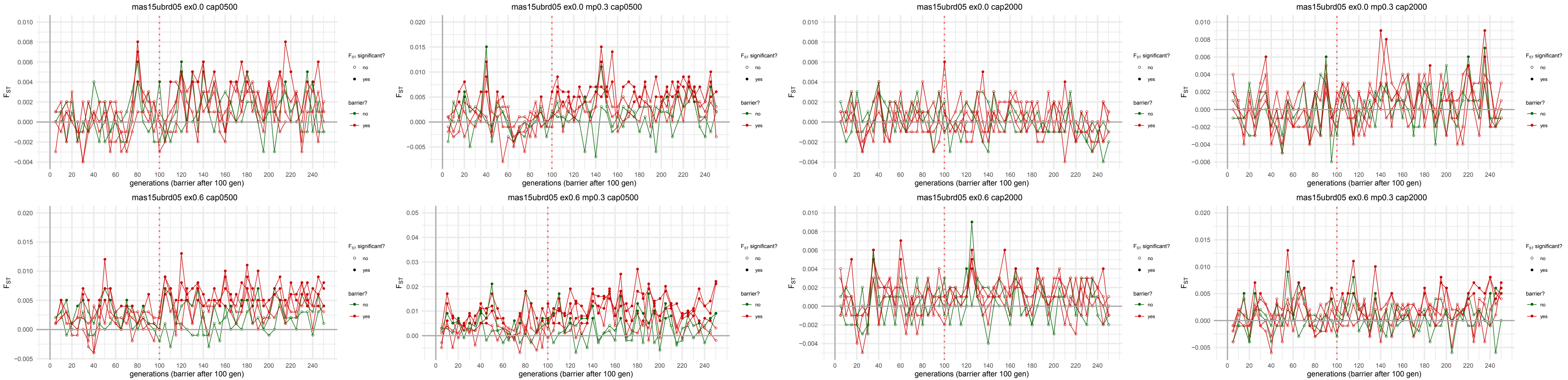
mas15ubrd05 ex0.0 cap0500
mas15ubrd05 ex0.0 mp0.3 cap0500
mas15ubrd05 ex0.0 cap2000
mas15ubrd05 ex0.0 mp0.3 cap2000
mas15ubrd05 ex0.6 cap0500
mas15ubrd05 ex0.6 mp0.3 cap0500
mas15ubrd05 ex0.6 cap2000
mas15ubrd05 ex0.6 mp0.3 cap2000
$F_{ST}$
generations (barrier after 100 gen)
$F_{ST}$ significant?
no
yes
barrier?
no
yes
