## Supplementary material for "Individual small in-stream barriers contribute little to strong local population genetic structure five strictly aquatic macroinvertebrate taxa": Figure S7

**Figure S7**: Plots showing changes in allelic richness (page 1 -2) and observed heterozygosity (page 3-4) among populations over generations for different Nemo simulation scenarios. The barrier, introduced after 100 generations is indicated by a red line. A representative subset of simulations is shown. Results were similar for all migration matrices and barrier models and are therefore only shown for *asymmetric15* (mas15) and the most differing barrier models (cb = complete barrier, ub = upstream barrier). Simulations without extiction (ex0.0) and an extinction rate of 60% per generation (ex0.6) are shown either for random mating populations (page 1 & 3) or for hermaphrodites (indicated by mp0.3 for mating proportion/selfing rate, page 2 and 4). Patch capacities of 100, 500 and 2000 are shown (cap0100/0500/2000).

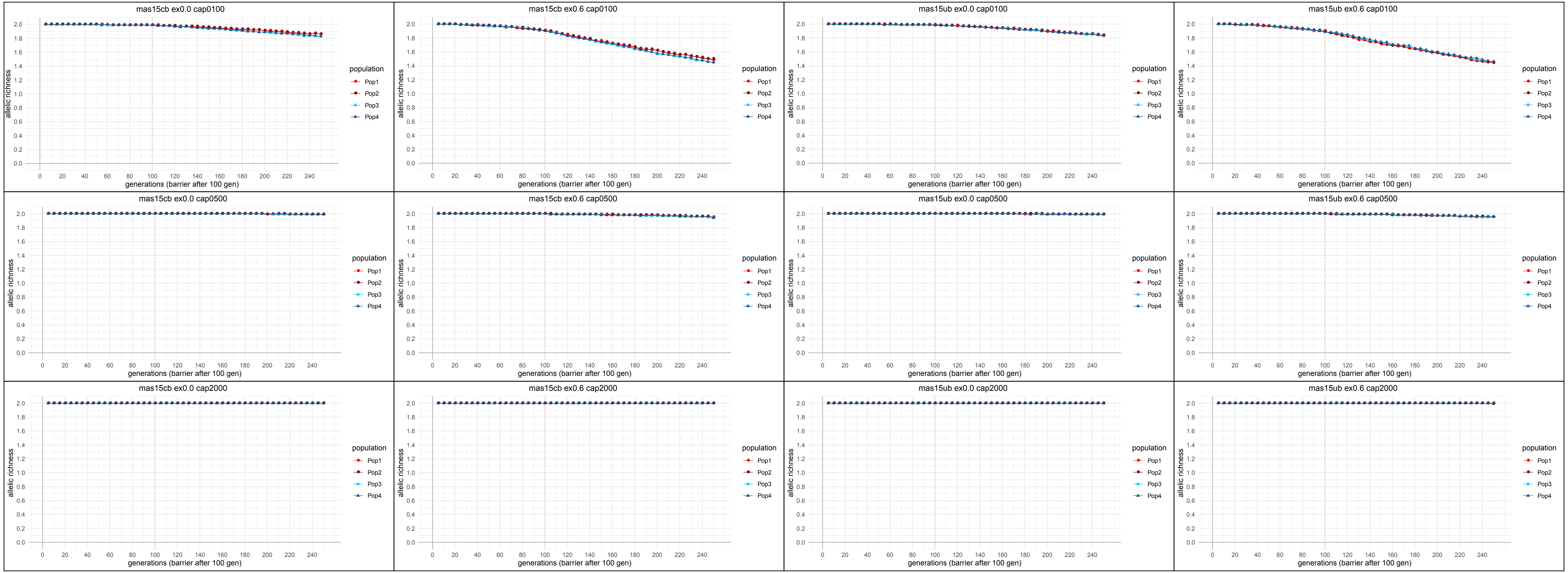

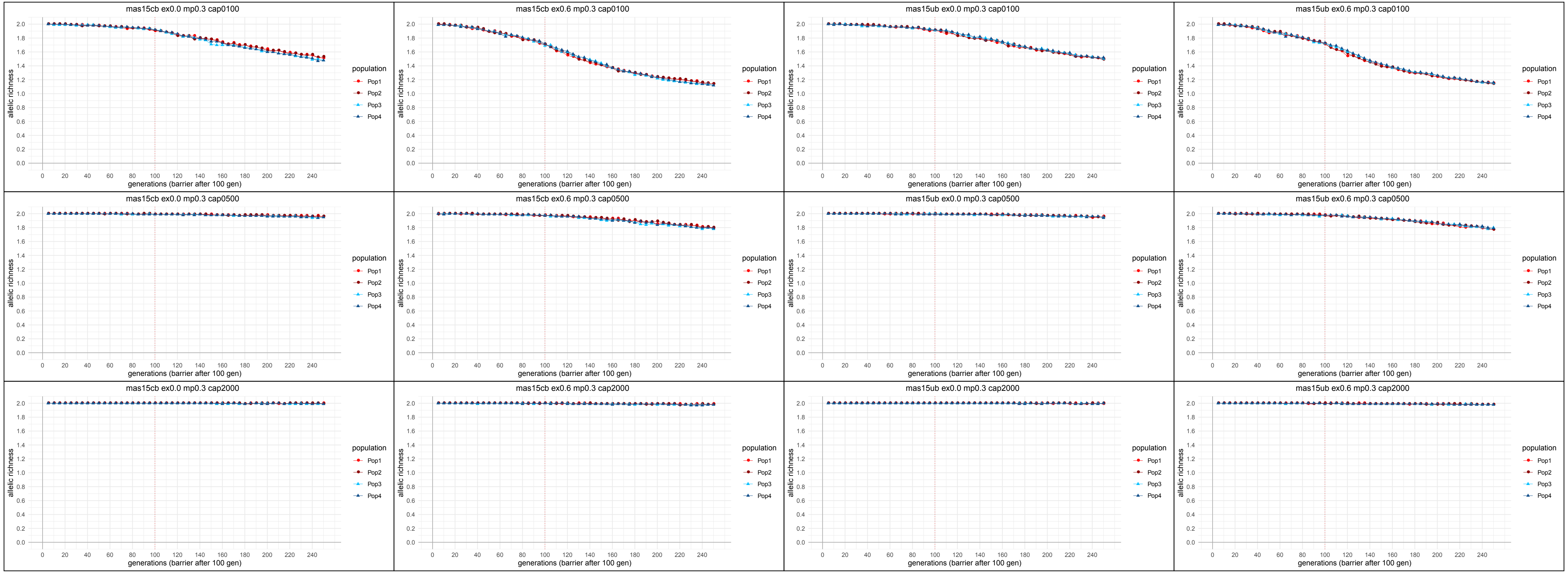
mas15cb ex0.0 mp0.3 cap0100
mas15cb ex0.6 mp0.3 cap0100
mas15ub ex0.0 mp0.3 cap0100
mas15ub ex0.6 mp0.3 cap0100
mas15cb ex0.0 mp0.3 cap0500
mas15cb ex0.6 mp0.3 cap0500
mas15ub ex0.0 mp0.3 cap0500
mas15ub ex0.6 mp0.3 cap0500
mas15cb ex0.0 mp0.3 cap2000
mas15cb ex0.6 mp0.3 cap2000
mas15ub ex0.0 mp0.3 cap2000
mas15ub ex0.6 mp0.3 cap2000
allelic richness
0.0
0.2
0.4
0.6
0.8
1.0
1.2
1.4
1.6
1.8
2.0
0
20
40
60
80
100
120
140
160
180
200
220
240
generations (barrier after 100 gen)
population
Pop1
Pop2
Pop3
Pop4

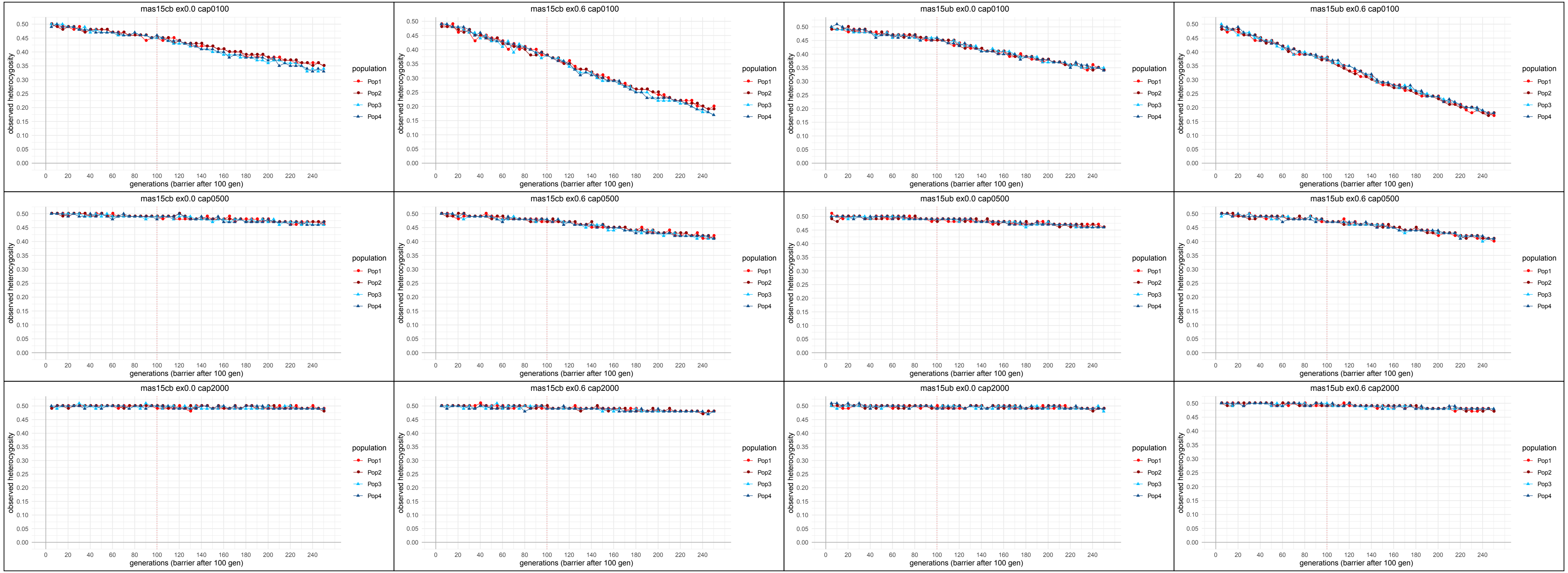
mas15cb ex0.0 cap0100
mas15cb ex0.6 cap0100
mas15ub ex0.0 cap0100
mas15ub ex0.6 cap0100
mas15cb ex0.0 cap0500
mas15cb ex0.6 cap0500
mas15ub ex0.0 cap0500
mas15ub ex0.6 cap0500
mas15cb ex0.0 cap2000
mas15cb ex0.6 cap2000
mas15ub ex0.0 cap2000
mas15ub ex0.6 cap2000
observed heterocygosity
generations (barrier after 100 gen)
population
Pop1
Pop2
Pop3
Pop4

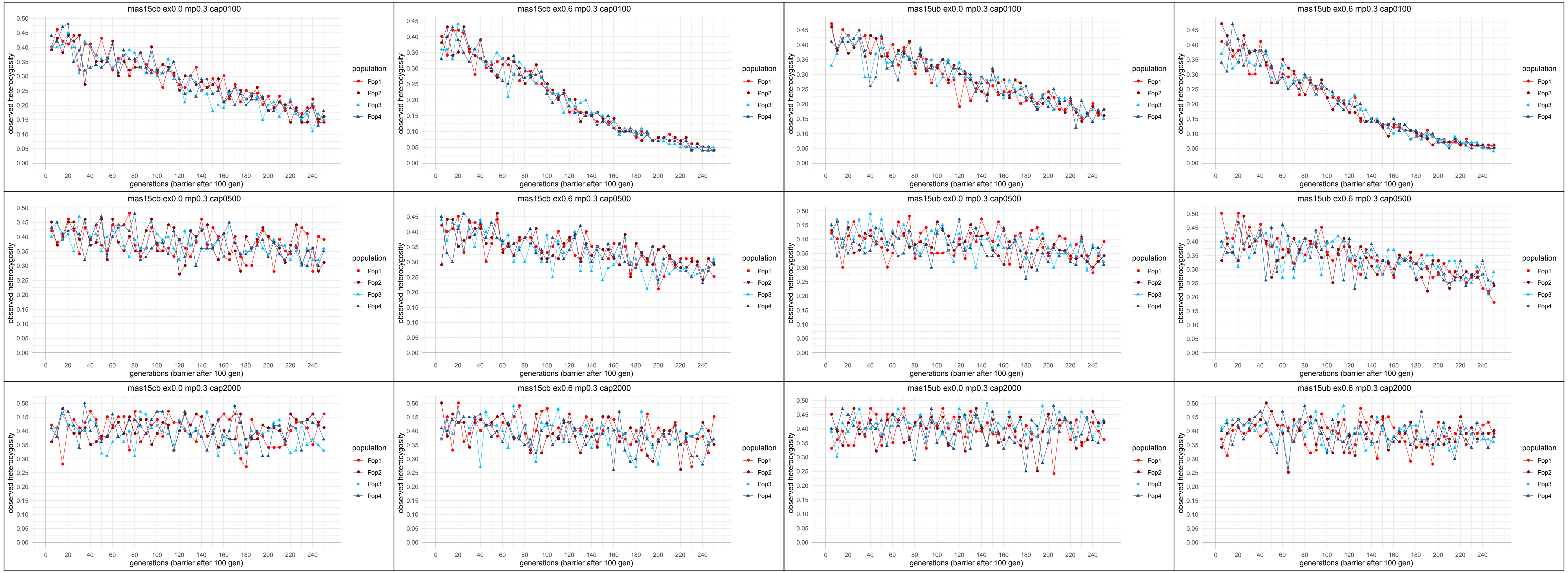

mas15cb ex0.0 mp0.3 cap0100
mas15cb ex0.6 mp0.3 cap0100
mas15ub ex0.0 mp0.3 cap0100
mas15ub ex0.6 mp0.3 cap0100
mas15cb ex0.0 mp0.3 cap0500
mas15cb ex0.6 mp0.3 cap0500
mas15ub ex0.0 mp0.3 cap0500
mas15ub ex0.6 mp0.3 cap0500
mas15cb ex0.0 mp0.3 cap2000
mas15cb ex0.6 mp0.3 cap2000
mas15ub ex0.0 mp0.3 cap2000
mas15ub ex0.6 mp0.3 cap2000
observed heterozygosity
generations (barrier after 100 gen)
population
Pop1
Pop2
Pop3
Pop4
